# Effects of ketamine and propofol on muscarinic plateau potentials in rat neocortical pyramidal cells

**DOI:** 10.1101/2024.02.14.579884

**Authors:** Anne S. Fleiner, Daniel Kolnier, Nicholas Hagger-Vaughan, Johan Ræder, Johan F. Storm

## Abstract

Propofol and ketamine are widely used general anaesthetics, but have different effects on consciousness: propofol gives a deeply unconscious state, with little or no dream reports, whereas vivid dreams are often reported after ketamine anaesthesia.

Ketamine is an N-methyl-D-aspartate (NMDA) receptor antagonist, while propofol is a γ-aminobutyric-acid (GABA_A_) agonist, but these mechanisms do not fully explain how these drugs alter consciousness.

Most previous *in vitro* studies of cellular mechanisms of anaesthetics have used brain slices or neurons in a nearly “comatose” state, because no “arousing” neuromodulators were added. Here we tested mechanisms of anaesthetics in slices after adding the cholinergic agonist muscarine to partly mimic an “awake-like” state.

Using whole-cell patch-clamp recordings from layer 2/3 pyramidal cells (L2/3PCs) in rat medial prefrontal cortex (mPFC) slices, we saw that muscarine induced long-lasting depolarizing plateau potentials (PPs) and spiking following brief depolarizing current injections. According to leading theories of consciousness and working memory, L2/3PCs and PPs are particularly important for these cognitive functions. After 2 hours of pre-incubation with ketamine or propofol, the muscarine-induced PPs were altered in different ways: 3 µM propofol reduced the PPs and (significantly) spiking, whereas 20 µM ketamine seemed to enhance PPs and spiking (non-significantly). Brief wash-in of these drug concentrations failed to induce such effects, probably due to insufficient equilibration by diffusion in the slices. In contrast, pre-incubation with 100 µM ketamine suppressed the PPs and spiking.

The different effects on PPs may be related to contrasting clinical effects: ketamine causing atypical anaesthesia with vivid, “psychedelic” dreaming while propofol causes less dreaming. However, high ketamine or propofol concentrations both suppressed PPs, suggesting possible connections between PPs, desynchronized activity, and consciousness. More experiments are needed to test these tentative conclusions.

## Introduction

Propofol and ketamine are both widely used for general anaesthesia (GA) in surgery, but have quite different effects on the state of consciousness (Franks 2008; G. Smith et al. 2023). Like other general anaesthetics they both induce unresponsiveness, as required for surgery (Franks 2008).

Such “unresponsiveness”, characterised by no overt response to or recall of pain or other stimuli, corresponds to the practical, clinical concept of “unconsciousness” that is commonly used in anaesthesiology and other clinical disciplines. This is as opposed to the concept of “unconsciousness” commonly used in consciousness research, where all forms of subjective experience, including “inner” experiences like dreaming, are regarded as conscious (Sanders et al. 2012). Recognizing this conceptual distinction, we here use the term “unresponsiveness” rather than “unconsciousness” for the mental state that is typical of general anaesthesia. However, it is important to note that the level of anaesthesia needed for unresponsiveness to severe pain for most anaesthetics except ketamine, is usually much deeper than the levels required for unresponsiveness to verbal commands or milder stimuli.

Propofol and ketamine also have quite different effects on “consciousness” in the latter sense of this term (i.e. all experience is defined as “conscious”). At surgical GA doses, propofol gives an unconscious state, with little or no dream reports after awakening (Nilsen, Arena, and Storm 2023; Nordström et al. 1997). In contrast, rich and vivid dreams, often intense and with “psychedelic” content, are frequently reported after surgical ketamine anaesthesia (Collier 1972; Grace 2003).

Most previous *in vitro* studies of cellular mechanisms of anaesthetics have used brain slices and neurons in a nearly “comatose” state, since no “arousing” neuromodulators, which are normally required for a physiological awake state, were added (Hoerbelt and Heifets 2018; Potez and Larkum 2008; Kitamura et al. 2003; Pittson, Himmel, and MacIver 2004; Yin et al. 2019; Steriade 2001). This is common practice in brain slice experiments (Lee et al. 2005; Dingledine 1984) although it is well known that depriving the cerebral cortex of arousing neuromodulatory input from the ascending reticular activation system (Garcia-Rill et al. 2013; Moruzzi and Magoun 1949), basal forebrain, and thalamus e.g. by cutting ascending ARAS axons, invariably causes coma *in vivo* (Posner et al. 2007; Timofeev et al. 2000). Of course, such a deprivation necessarily happens when making isolated cortical slices (Steriade 2001; Sanchez-Vives, Massimini, and Mattia 2017). However, in our opinion, this commonly used *in vitro* approach, starting with isolated brain tissue that is already in a state corresponding to “coma”/“unconsciousness” of the intact brain *in vivo*, seems to make little sense when studying mechanisms of how anaesthetics induce unconsciousness, since the contrast in states that one wants to study (“conscious” vs. “unconscious” states) is already precluded before the experiments start.

Therefore, in this study, we decided to test mechanisms of anaesthetics *in vitro* after first trying to induce or mimic a partly “awake-like” state in the cortical tissue, by bath-applying the cholinergic agonist muscarine before testing the effects of the anaesthetics. We chose to use only muscarine for this purpose, because the muscarinic effects of ACh on cortical pyramidal cells are particularly powerful (McCormick and Prince 1985; Nuñez et al. 2012; Williams and Fletcher 2019), and can be induced in an easily controllable manner. Ideally, one would like to closely mimic an “awake-like” state in slices by supplying a cocktail of all the relevant neurochemicals at physiological concentrations, and apply them in the normal temporo-spatial patterns that the cortical tissue is normally exposed to in the awake, behaving animal. However, as this is currently impossible to achieve *in vitro*, we opted for a highly simplified, easily controllable approach, using only a single agonist, muscarine. An additional reason for this choice, is that we have recently found that muscarine alone induces plateau potentials in layer 2 and 3 pyramidal cells (L2/3PCs) in rat medial prefrontal cortex (mPFC) - a mechanism that profoundly alters the input-output function of the neurons, and thus may be important for transitions between conscious and unconscious states, and hence possibly for anaesthesia (Hagger-Vaughan, Kolnier, and Storm 2023).

Another type of problem with most *in vitro* studies of anaesthetics (in addition to “comatose” brain slices), is the difficulty of relating clinically relevant concentrations and effects of anaesthetic drugs in humans during general anaesthesia, to drug levels used and effects observed in isolated brain slices lacking blood supply. We get back to these complex issues in the Discussion. In this study, we used concentrations of ketamine (20 µM) and propofol (3 µM) that seem likely to approach (albeit higher than) the estimated clinically relevant free concentrations of these drugs in the human brain, but lower than concentrations often used in previous brain slice studies. In addition, we tested higher concentrations for comparison (100 µM ketamine; and 10 µM propofol).

The cellular and molecular mechanisms underlying clinical GA with ketamine and propofol are also known to be very different (Franks 2008), but still remain to be fully elucidated. Furthermore, the connections between the mechanisms of these two anaesthetics and their different effects on consciousness (reported subjective experience) also remain to be fully understood (Franks 2008).

Like most general anaesthetics, propofol is a GABA_A_ receptor potentiator (Edwards and Preuss 2023; Trapani et al. 2000) (positive allosteric modulator) leading to enhanced inhibition in the brain, which is supposed to lead to sedation and loss of consciousness (Franks 2008). In contrast, the anaesthetic effect of ketamine is mainly ascribed to its noncompetitive N-methyl-D-aspartate (NMDA) receptor antagonism, leading to reduced excitatory neurotransmission and unresponsiveness (Franks 2008). Another possibility is that ketamine, acting on different NMDAR subtypes, partly blocks NMDARs that stimulate inhibitory GABAergic interneurons, thus causing disinhibition and increased network activity (McMillan & Muthukumaraswamy, 2020). However, these different mechanisms do not seem to entirely explain how these drugs alter consciousness and responsiveness.

Propofol is currently the most common anaesthetic used to induce and maintain surgical anaesthesia (Hofschneider 2018). In addition to its potentiation of GABA_A_ receptors, propofol has also been found to affect a number of other molecular targets; e.g. it inhibits hyperpolarization-activated cyclic nucleotide–gated (HCN) channels (Ying et al. 2006), calcium-activated potassium channels (Ying and Goldstein 2005), L-type voltage-gated calcium channels (Fassl et al. 2011), and transient receptor potential (TRP) channels (Bahnasi et al. 2008).

Ketamine is most commonly used as an anaesthetic in hypovolemic patients and trauma surgery due to its favourable cardiovascular effects in such patients (Kurdi, Theerth, and Deva 2014). More widespread use of ketamine is restricted mostly due to the risk of unpleasant, nightmare-like awakening experienced in many patients (White, Way, and Trevor 1982). Similarly to propofol, a wide range of other molecular targets for ketamine have been found (in addition to NMDA receptors), which could also contribute to the anaesthetic effect, including inhibition of HCN channels (Chen, Shu, and Bayliss 2009), acetylcholine release (Lydic and Baghdoyan 2002), and L-type calcium channels (Yamakage, Hirshman, and Croxton 1995), and enhancement of BDNF signalling (Moliner et al. 2023).

Studies using transcranial magnetic stimulation (TMS) combined with EEG to assess consciousness in humans (the perturbational complexity index, PCI, method; (Casali et al. 2013) found that the brain (cortex) is in a so-called “low-complexity state” during propofol anaesthesia (Sarasso et al. 2015) with an EEG pattern modestly resembling that of sleep (Casey et al. 2022), and there were few or no post-anaesthesia reports of dreaming (Sarasso et al. 2015). In contrast, ketamine anaesthesia leads to a state of unresponsiveness with a high-complexity brain state (Sarasso et al. 2015) with EEG patterns resembling those of wakefulness (Colombo et al. 2019), and is often followed by recollections of vivid dreams experienced during anaesthesia (Collier 1972; Grace 2003).

Since reports of dreams or other conscious experiences are also so much less common following propofol compared to ketamine anaesthesia (Nordström et al. 1997), this has led to the interpretation that whilst both drugs lead to unresponsiveness to pain and other stimuli (i.e, surgical anaesthesia), a form of consciousness is retained during ketamine, leading to internal experience without responsiveness, similar to that during REM sleep (Aru et al. 2020; Hobson 2009). To help elucidate the causes of these differences, it is important to further investigate the cellular effects of ketamine in comparison with more typical anaesthetics such as propofol.

The “seat” of consciousness in the brain is a highly contested topic (Boly et al. 2017). The prefrontal cortex (PFC) is implicated as a key area in several leading theories including the global neuronal workspace theory (Mashour et al. 2020) and higher-order thought (HoT) theories (Brown, Lau, and LeDoux 2019), as well as in functions related to consciousness such as attention (Moore and Armstrong 2003; Clark et al. 2015), working memory (Asaad, Rainer, and Miller 1998), and emotional processing (Burgos-Robles et al. 2017). Therefore, this area is of great interest for the study of effects of anaesthetics on the physiology of individual neurons.

Within the PFC, layer 2/3 pyramidal cells (L2/3PCs) are particularly relevant for the “frontal” theories of consciousness (GNWT; HOT) due to their long-range axonal projections, which make extensive corticocortical and subcortical connections that support the recurrent processing required by these theories (Wang 2001). Thus, the L2/3PCs with their long axons are supposed to form the main basis for the “global workspace” of the GNWT (Dehaene, Kerszberg, and Changeux 1998; Goldman-Rakic 1988; Mashour et al. 2020).

The ability to switch to conscious states is mainly caused by long ascending projections from the reticular activation system (ARAS) and basal forebrain nuclei, which provide neuromodulatory input to the cortex, including acetylcholine (ACh) and monoamine neurotransmitters (Paus 2000). The combined effect of neuromodulator release following ARAS activation is a transition from sleep to wakefulness (Steriade, Timofeev, and Grenier 2001), and wakefulness and conscious states are associated with high cortical levels of acetylcholine and monoamines (Watson, Baghdoyan, and Lydic 2010; Lee et al. 2005). When the modulatory input from the ARAS and basal forebrain is severely reduced, for example by brain stem lesions, with consequent loss of its long-range cholinergic and monoaminergic projections to the cortex and forebrain, the individual enters a comatose state, with loss of consciousness and normal, awake activity (Edlow et al. 2020). Therefore the effects of these neuromodulators on neuronal and synaptic physiology are of great interest.

ACh can induce broad, lasting changes in neuronal physiology via the signalling cascades induced by activation of muscarinic ACh receptors (mAChRs). Muscarinic effects include drastic alterations of spike frequency adaptation, which is a key parameter for transition to the aroused state (Dasilva et al. 2021; Cole and Nicoll 1984; D’Andola et al. 2018). ACh can also induce powerful plateau potentials (PPs) (Fraser and MacVicar 1996; Egorov et al. 2002; Williams and Fletcher 2019), i.e. extended periods of depolarization and action potential firing which often greatly outlast the initial depolarizing stimulus, and fundamentally alters the input-output functions of these neurons in the awake state compared to unconscious states.

PPs have been implicated in the induction of long-term potentiation (Gambino et al. 2014), a likely mechanism of long-term memory (Bliss and Collingridge 1993), and working memory (Egorov et al. 2002), by keeping information present in a network beyond the termination of a sensory stimulus (Mashour et al. 2020). PPs have been suggested to be fundamentally involved in consciousness by acting as a mechanism of detecting coincident sensory and contextual input and amplifying action potential output, allowing that information to enter conscious awareness (Aru, Suzuki, and Larkum 2021). PPs have been observed in a range of different neuron types and brain areas (Hagger-Vaughan and Storm 2019; Lo and Erzurumlu 2002; Fraser and MacVicar 1996; Oikonomou et al. 2014). Recently we found that muscarine alone induces PPs in PFC L2/3PCs following depolarizing current injections, and characterised the molecular and morphological underpinnings of these PPs (Hagger-Vaughan, Kolnier, and Storm 2023).

As PPs may be a key mechanism for supporting consciousness, their sensitivity to anaesthetics is of interest, in particular whether the effects of ketamine and propofol on PPs differ, which might help explain their very different effects on brain state and consciousness.

In this *in vitro* patch clamp study we investigate and compare the effects of brief and extended exposure to a range of relevant concentrations of ketamine or propofol on muscarinic PPs in L2/3PCs in the PFC. We find that brief exposure to high concentrations of ketamine (100 µM) or propofol (10 µM), and long incubation with lower concentrations of propofol (3 µM) caused a significantly reduced incidence (and reduced amplitude for 100 µM ketamine) of PPs, with less associated spiking. We discuss whether and how these cellular effects may contribute to the different anaesthetic effects of ketamine and propofol.

## Materials and Methods

### Ethical Approval

All animal procedures were approved by the responsible veterinarian of the institute, in accordance with the statute regulating animal experimentation (Norwegian Ministry of Agriculture, 1996).

### mPFC slice preparation, maintenance, and perfusion of recording chamber

Coronal prefrontal slices were obtained from young (P21-P28), male Wistar rats (Scanbur AS, Nittedal, Norway). Rats were anaesthetised with isoflurane inhalation and decapitated. The brain was removed quickly into ice-cold sucrose-based artificial cerebrospinal fluid (aCSF) containing (in mM): 87 NaCl, 1.25 KCl, 1.25 KH_2_PO_4_, 7 MgCl_2_, 0.5 CaCl_2_, 16 glucose, 75 sucrose, 25 NaHCO_3_) saturated with 95% O_2_–5% CO_2_. 400 μm slices were cut using a Leica VT1200 vibratome (Leica Microsystems; Wetzlar, Germany) and incubated for 30 min at 35°C in aCSF containing (in mM): 125 NaCl, 2.5 KCl, 1.25 NaH_2_PO_4_, 1.4 MgCl_2_, 1.6 CaCl_2_, 16 glucose, 25 NaHCO_3_) saturated with 95% O_2_–5% CO_2_. After incubation slices were kept at room temperature (∼20°C) until use.

For propofol experiments, PTFE tubing was used for delivering aCSF to the slice chamber to avoid absorption of propofol into silicone or Tygon tubing, as has previously been reported (Maurer et al. 2017). Propofol was freshly diluted on the day of the experiments. For experiments using slices that were pre-incubated with either ketamine or propofol, we used glass holding chambers for the pre-incubation to avoid absorption of any drugs, thereby diminishing the effective concentration in the medium. These slices were incubated for at least two hours before performing recordings, to allow the drugs to equilibrate in the tissue. It was previously shown that 2 hours incubation is sufficient to reach concentrations close to equilibrium for propofol near the surface of the slices, at depths down to ∼50 µm, where all our somatic whole-cell recordings were obtained, and 2 hours far exceeded the time taken to reach equilibrium for ketamine (Gredell et al. 2004; Geiger et al. 2021). Control cells in these experiments were from slices incubated for the same duration in standard aCSF, to compensate for any potential time-dependent effects of the incubation per se. In some experiments, the synaptic blockers SR 95531 (gabazine; 5 µM), DNQX (10 µM), and AP5 (50 µM) were included in the aCSF to block glutamatergic and GABA_A_-ergic transmission. No differences in the amplitude, duration, or induced spiking of plateau potentials were observed between experiments with or without synaptic blockers, therefore data from both conditions have been merged for the subsequent analysis.

### Electrophysiology

Whole-cell current-clamp recordings were obtained using visual guidance from IR-DIC optics (BX51WI; Olympus, Tokyo, Japan) from the somata of L2/3 mPFC pyramidal cells. Slices were maintained at 32 ± 1°C and superfused with aCSF. Patch-clamp pipettes (5–7 MΩ) were pulled from borosilicate glass tubing (outer diameter 1.5 mm, inner diameter 0.86 mm, with filament; Sutter Instruments, Novato, CA, USA) and filled with a solution containing (in mM): 120 K-methanesulfonate, 4 Na_2_ATP, 0.3 NaGTP, 10 Na_2_phosphocreatine, 10 KCl, 3 MgCl_2_, 10 inositol, 10 HEPES. The pH of the intracellular medium was adjusted to 7.2-7.3 with KOH, and osmolarity was between 290 and 300 mOsmol^−1^. Recordings were made using either a Dagan BVC-700A patch-clamp amplifier (Dagan Corporation, Minneapolis, MN, USA) or a Multiclamp 700A (Molecular Devices, San Jose, CA, USA), low-pass filtered at 10 kHz, and digitised at 20 kHz. Access resistance was typically between 10 and 30 MΩ and was compensated at the beginning of every recording and adjusted as required.

### Stimulation protocol

Injecting trains of short (2 ms), depolarizing current pulses (1.0-1.5 nA; amplitude adjusted to elicit only a single spike per pulse), we repeatedly induced trains of 7 action potentials at 70 Hz, once every 5 minutes.

### Data Acquisition and Analysis

Data were acquired using pCLAMP 10 software and digitised with either a Digidata 1322A or Digidata 1440 (Molecular Devices). The analysis was carried out using Clampfit software (Molecular Devices), and results were plotted and statistical analysis performed in Origin 2020 (OriginLab Corp; Northampton, MA, USA) and GraphPad Prism, version 9. Cells were manually held at a baseline membrane potential of −60 mV by DC current injections. Recordings were made in current-clamp after obtaining a seal of 1 GΩ or tighter. The potentials reported here were not corrected for the junction potential, which was calculated to be 9.4 mV between the intracellular solution and the aCSF used here. The plateau potential was quantified using two different parameters: the post-burst area and post-burst spikes. The post-burst area was defined as the area under the curve (either positive-going or negative-going) in the first 10 seconds following the initial elicited spike burst (**Figure 1B**; area indicated by grey shading). Post-burst spikes were defined as the number of spikes observed in the first 10 seconds following the initial elicited spike burst (**Figure 1B** lower trace and inset). In wash-in experiments, we excluded all cells that did not elicit spiking plateau potentials after the wash-in of muscarine to increase the probability that we were testing a quite uniform subpopulation of cells.

**Figure 1.**
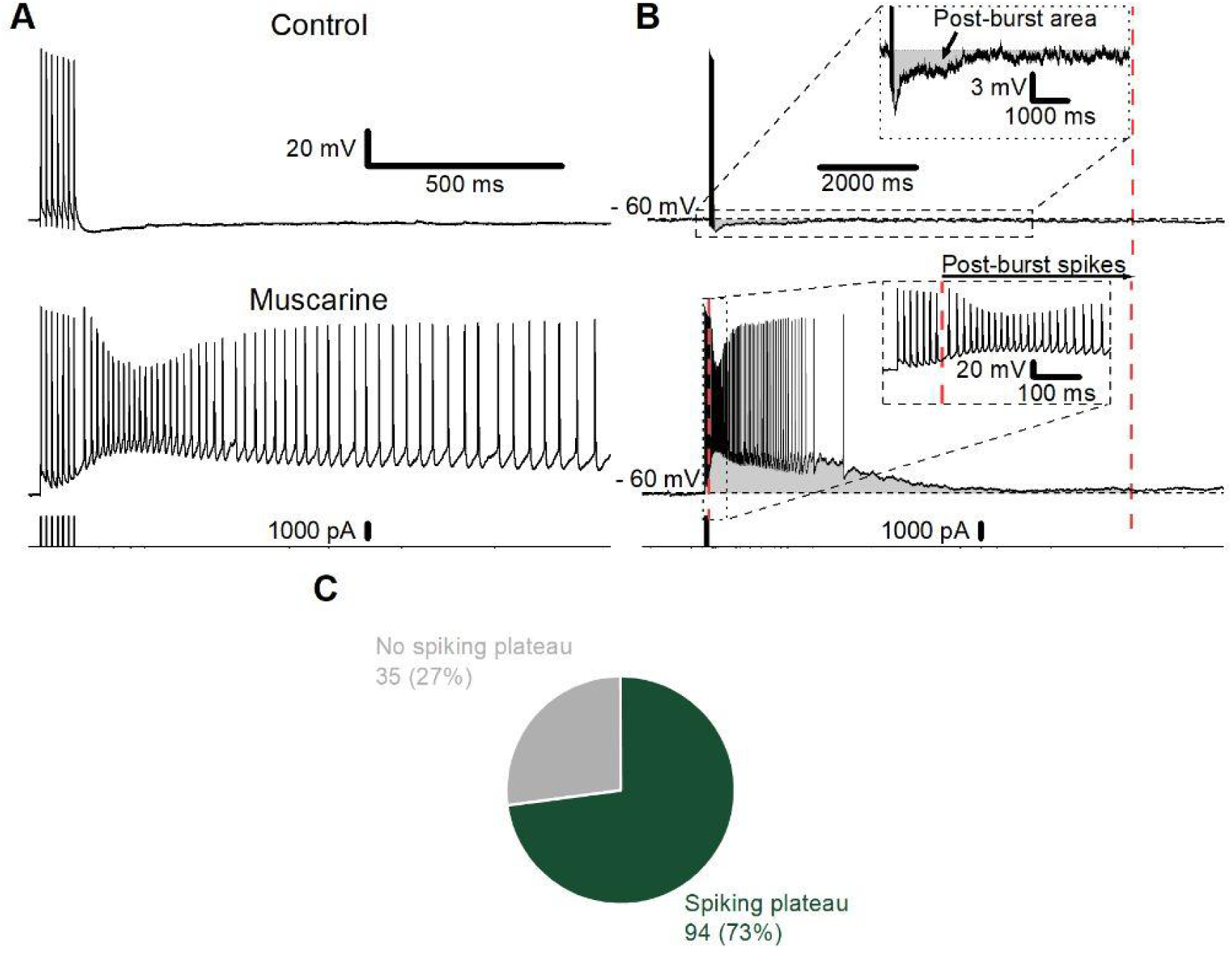
Experimental details and analysis of muscarine-induced plateau potentials. A) Example traces of the membrane potential following 7 action potentials at 70 Hz elicited by the injection of 7 brief, depolarizing current pulses (lower trace) in control conditions (top) and following bath application of 10 µM muscarine (middle). B) Typical examples of traces before (top) and after muscarine application (bottom) to show how plateau potentials (PPs) were measured. The area of the after-potentials, i.e. the membrane potential deviation after the 7th spike (the deviation from the mean membrane potential before the 1st spike) is marked by grey shading. In the absence of muscarine (Control; upper trace; enhanced scale inset), there was an afterhyperpolarization (AHP) after the spikes, whereas the muscarine-treated cells showed a long-lasting afterdepolarization, called a “plateau potential” (PP, lower trace). The number of action potentials following the initial burst is referred to as ‘spikes’ and is illustrated in the inset of the lower trace. The area measurements and spike counts started immediately after the spikes elicited by current injections. The red, dotted lines on the far-right of the traces denote the endpoints of the measurements and spike counts. C) Summary chart showing the proportion of cells (%) exhibiting a spiking plateau following the application of muscarine. Only cells from slices incubated in control aCSF (no ketamine or propofol) are included. Raw cell count is shown adjacent to each sector.

Data are reported as the mean ± standard error of the mean (SEM). Non-parametric tests were used to compare groups as group sizes tended to be too small to determine whether they were normally distributed. Statistical comparisons were made using Wilcoxon-signed rank test for paired data and Mann-Whitney U test for unpaired data. Normalising of data was performed by dividing the value at each time point in the individual trials by the area or number of spikes after the wash-in of muscarine (time = 5 minutes). We normalised the data from wash-in experiments (**Figures 2** and **4**) to investigate the relative change in area and spiking after the wash-in of propofol and ketamine. The data from the pre-incubation experiments (**Figures 3** and **5**) was not normalised, as the absolute area and spiking value are of interest.

**Figure 2.**
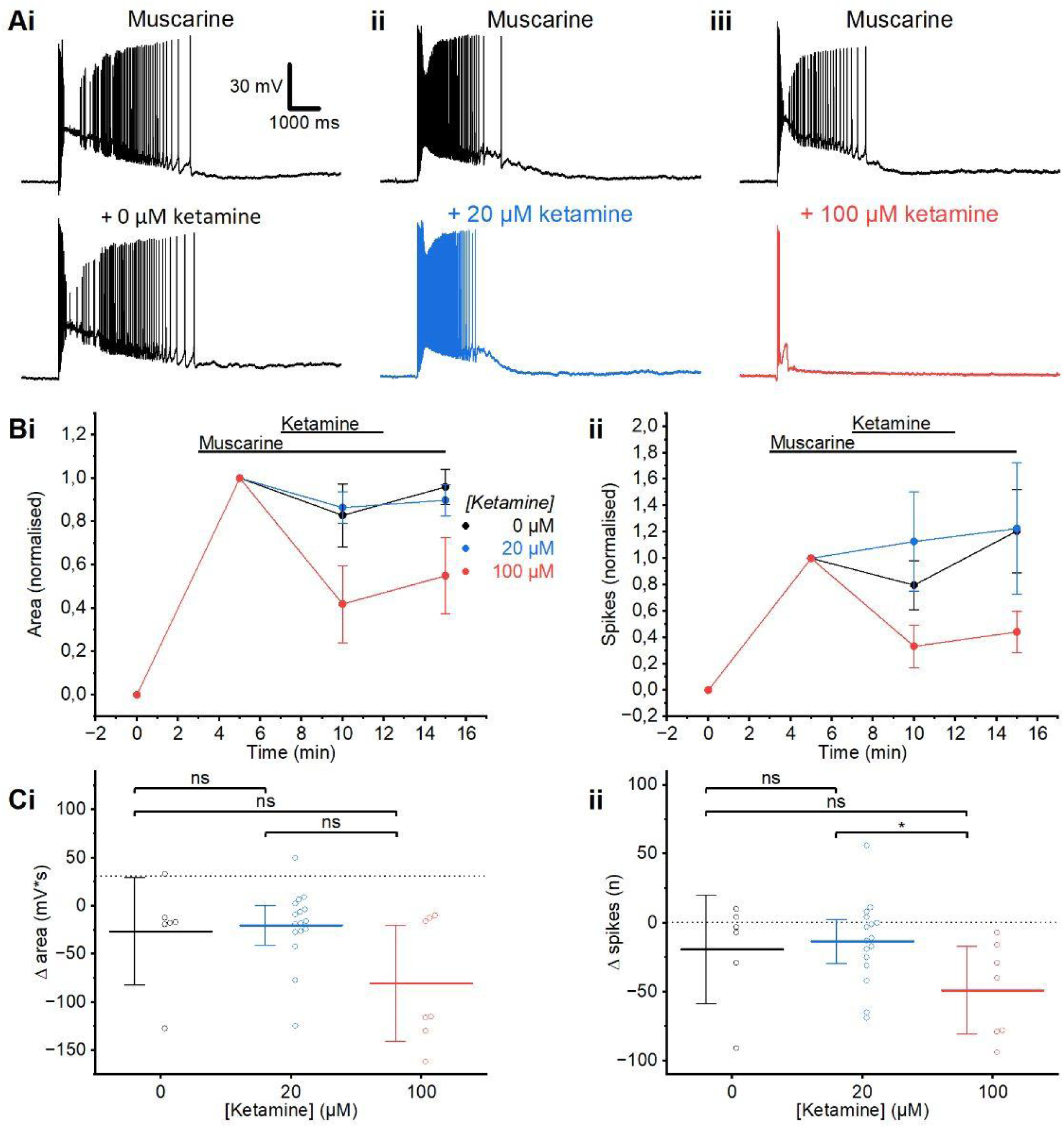
Ketamine wash-in. A) Example traces showing the effect on the muscarinic plateau of the 5-minute wash-in of ketamine (20 μM (ii) or 100 μM (iii) compared to the time-matched control, where the whole-cell recordings were maintained for the same period of time (5 min), but without adding any ketamine (0 μM (i), ketamine). B) Time course plots showing the normalised area (i) and spikes (ii) following the wash-in of muscarine and then the subsequent wash-in, then wash-out of ketamine at different concentrations (0 µM: n=6; 20 µM: n=16; 100 µM; n=7; Error bars = SEM). We plotted normalised values here, to visualise the ketamine effects relative to the different plateau sizes and spike counts before adding ketamine (see black traces in Ai-Aiii). C) Summary plots of the change in area (i) and spikes (ii) following application of ketamine at different concentrations, relative to prior to the wash-in of ketamine in the presence of muscarine. (i) The change in the post-burst area was not significantly different at any of the tested ketamine concentrations (MWU test, 0 µM; n=6, 20 µM; n=16, 100 µM; n=7). (ii) The change in post-burst spiking after the wash-in of 20 µM and 100 µM was not significant compared to the control (MWU test). However, the decrease in post-burst spiking was significantly larger after the wash-in of 100 µM ketamine than with 20 µM (MWU test). Error bars = 95% CI. * = p<0.05, ns = p>0.05.

**Figure 3.**
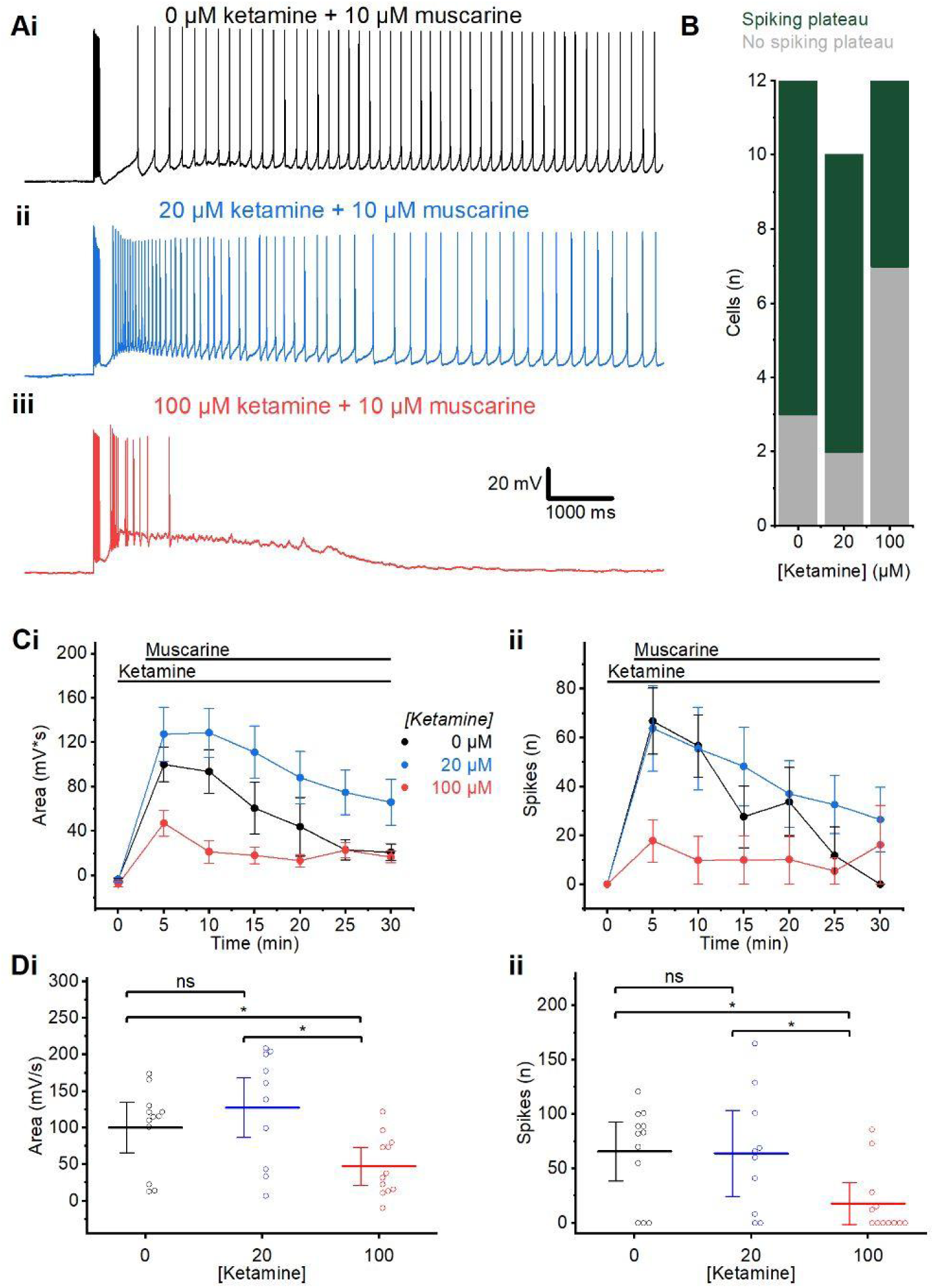
Ketamine pre-incubation. A) Example traces showing plateau potentials induced by 10 µM muscarine, following long-lasting (120 minutes) pre-incubation without muscarine, and without ketamine (0 µM) (i), or with 20 µM (ii), or 100 µM (iii) ketamine. B) Summary chart showing the number of cells exhibiting a spiking plateau or no spiking plateau, following incubation in control aCSF, 20 µM, and 100 µM ketamine. C) Time course plots showing the area (i) and spikes (ii) following the wash-in of muscarine in the presence of ketamine at different concentrations. Error bars = SEM. D) Summary plots of the area (i) and spikes (ii) following incubation in ketamine at different concentrations, measured at the first elicited spike burst after the wash-in of muscarine (time = 5 minutes). (i) No significant difference in post-burst area between the control (0 µM) and 20 µM ketamine was observed. The post-burst area was significantly smaller after wash-in of muscarine in slices incubated in 100 µM ketamine than in 0 µM and 20 µM (MWU test; 0 µM: n=12; 20 µM: n=10; 100 µM: n=12). (ii) No significant difference in spiking was observed between slices incubated in 0 µM and 20 µM ketamine, but there was a significant difference between 0 µM and 100 µM, and 20 µM and 100 µM ketamine (MWU test; 0 µM: n=12; 20 µM: n=10; 100 µM: n=12). Error bars = 95% CI. * = p<0.05, ns = p>0.05.

**Figure 4.**
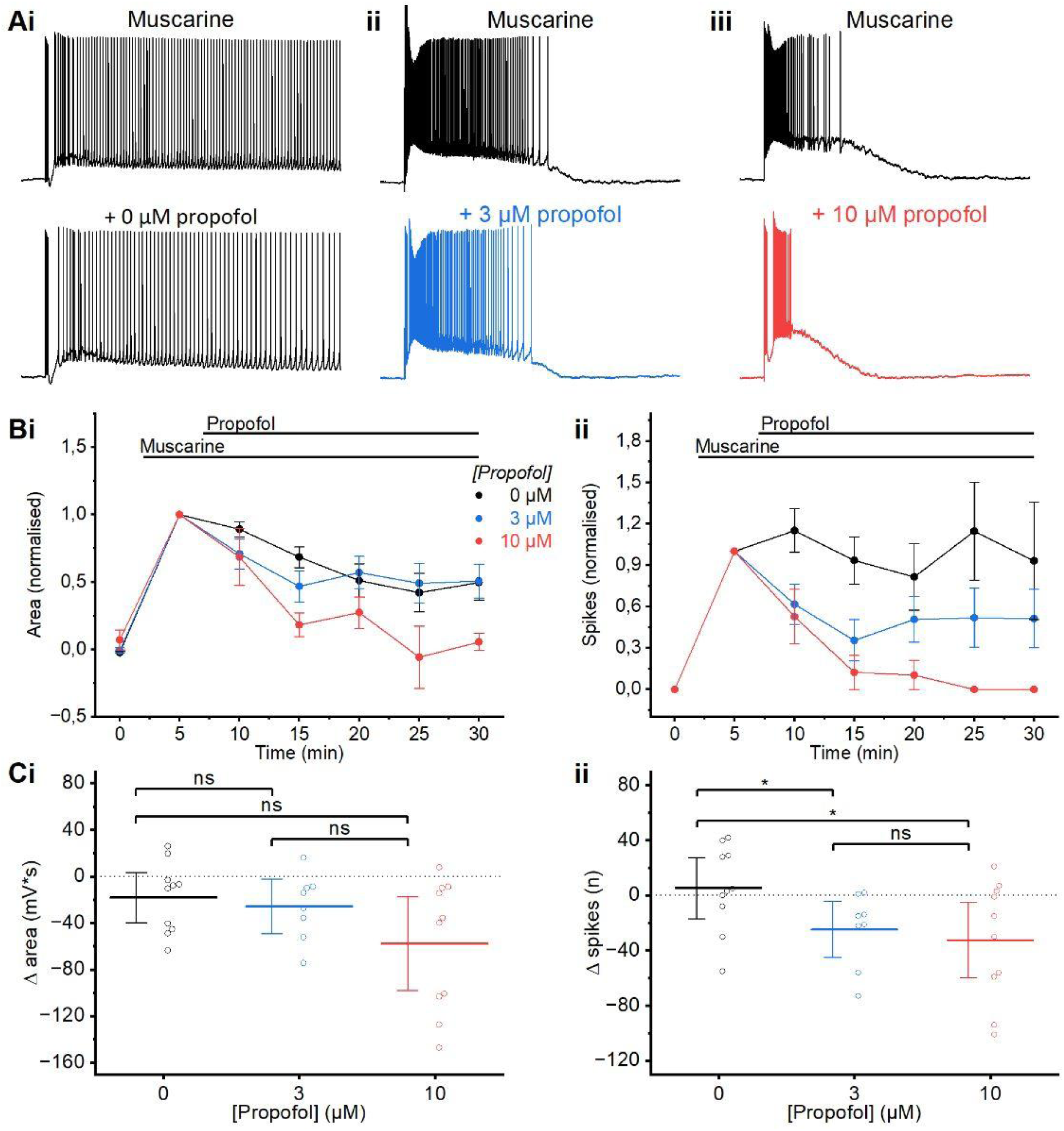
Propofol wash-in. A) Example traces of the muscarinic plateau before and after the wash-in of propofol 0 µM (i), 3 µM (ii) or 10 µM (iii). B) Time course plots showing the normalised area (i) and spikes (ii) following the application of propofol. Data are normalised to values observed following muscarine application, before application of propofol. Error bars = SEM. We plotted normalised values here, to visualise the propofol effects relative to the different plateau sizes and spike counts before adding propofol (see black traces in Ai-Aiii). C) Summary plots of the change in area (i) and spikes (ii) following application of propofol at different concentrations relative to prior to the wash-in of propofol (i) There was no difference in the change in the post-burst area between any of the concentrations of propofol used following 5 minutes of wash-in (MWU test; 0 µM: n=10; 3 µM: n=8; 10 µM: n=10). (ii) A significantly larger decrease in post-burst spikes was observed after wash-in of both 3 µM and 10 µM propofol compared to the control (0 µM). The decrease in spiking after wash-in of 3 µM and 10 µM was not significantly different (MWU test; 0 µM: n=10; 3 µM: n=8; 10 µM: n=10). Error bars = 95% CI. * = p<0.05, ns = p>0.05.

**Figure 5.**
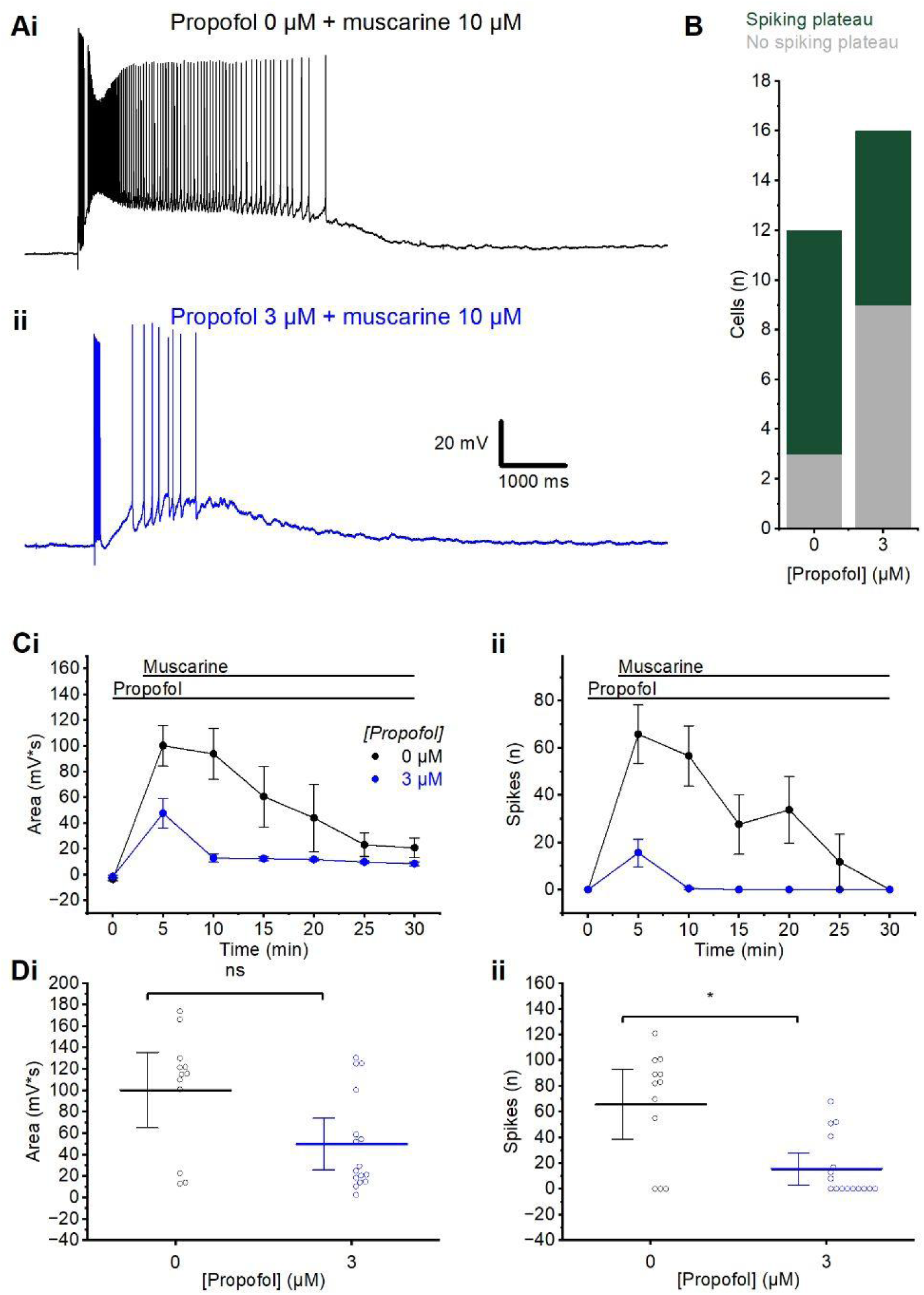
Propofol pre-incubation. A) Example traces of the muscarinic plateau potential in the presence of propofol 0 µM (i) or 3 µM (ii). B) Summary chart showing the proportion of cells exhibiting a spiking plateau or no spiking plateau, following incubation in control aCSF or 3 µM propofol. C) Time course plots showing the area (i) and spikes (ii) following the application of muscarine after slice incubation in propofol. Error bars = SEM. D) Summary plots of the change in area (i) and spikes (ii) following incubation in control aCSF or 3 µM propofol, measured at the first elicited spike burst after the wash-in of muscarine (time = 5 minutes). (i) Wash-in of muscarine after incubation in 3 µM propofol had no significantly different effect on post-burst area from incubation in 0 µM propofol (MWU test; 0 µM: n=12; 3 µM: n=16). (ii) There were significantly fewer post-burst spikes after wash-in of muscarine in slices incubated in 3 µM than in control-incubation (MWU test; 0 µM: n=12; 3 µM: n=16). Error bars = 95% CI. * = p<0.05, ns = p>0.05.

### Chemicals

Muscarine was obtained from Tocris Bioscience (Bristol, UK), racemic ketamine from Abcur AB (Helsingborg, Sweden), propofol from Sigma-Aldrich Norway AS (Oslo, Norway), DNQX and APV from Alemone (Jerusalem, Israel). Potassium gluconate, KMeSO_3_, and the other substances used for preparing the solutions were obtained from Sigma-Aldrich Norway AS (Oslo, Norway). All chemicals tested in the experiments were bath applied at a superfusion rate of ∼3 ml/min. The reasons for choosing the concentrations of ketamine and propofol that we used are discussed in the Introduction and Discussion.

## Results

### Muscarinic plateaus in the mPFC

We obtained stable somatic whole-cell recordings from L2/3 pyramidal cells (n=129) in brain slices from the medial prefrontal cortex (mPFC). A mean resting potential of −70.6±0.66 mV was observed under control conditions in these cells (n=40), with an input resistance of 96.3 ±5.25 MΩ (n=32). No spontaneous spiking was observed before or after adding muscarine in any of the cells under the conditions used here. Cells were manually held at a membrane potential of −60 mV by a positive holding current of 146±12.4 pA (n=45).

By injecting trains of short (2 ms), depolarizing current pulses (1.0-1.5 nA; amplitude adjusted to elicit only a single spike per pulse), we repeatedly induced trains of 7 action potentials at 70 Hz, once every 5 minutes. After bath application of muscarine (10 µM; **Figure 1A**), each spike train was followed by a plateau potential (PP), as described previously (Hagger-Vaughan, Kolnier, and Storm 2023). The PPs and spiking were quantified by measuring (1) the area under the PP curve (area; **Figure 1B** upper trace), and (2) the number of PP-triggered action potentials (spike number; **Figure 1B** lower trace) following the elicited burst of 7 spikes. The area was calculated as the integral of the membrane potential deviation from the baseline holding membrane potential in the 10 seconds following the offset of the current pulse train.

We have previously found that this concentration of muscarine most reliably induced plateau potentials in this cell type (Hagger-Vaughan, Kolnier, and Storm 2023), and here we observed this again, with 73% of cells exhibiting a spiking plateau, while 27% exhibited no spiking plateau (**Figure 1C**).

### Effects of ketamine on muscarinic plateau potentials

We then proceeded to test the effect of general anaesthetics on the plateau potential. Ketamine is noted for exerting a variety of clinical effects, as a psychedelic, an antidepressant, and an anaesthetic, depending on the dose (Mion and Villevieille 2013). It is also currently unclear what the effective concentration of ketamine is in the brain for a given intravenous or intraperitoneal dose (see discussion). For these reasons, we tested the effects of both 20 µM and 100 µM of ketamine, applied after induction of muscarinic plateaus.

We first used wash-in (bath-application) of ketamine. After wash-in of 20 µM ketamine **(Figure 2Aii** and **B**) we found no significant difference in the change in the plateau area (MWU; 0 µM: −26.6±21.7 mV*s, n=6; 20 µM: −20.3±9.6 mV*s, n=16, p=0.91) or spike number (MWU; 0 µM: −19±15 spikes, n=6; 20 µM: −14±7.5 spikes, n=16, p=0.96) (**Figure 2Ci-ii**).

However, we found that the mean number of spikes triggered by each PP was reduced, although not significantly, following the wash-in of a higher concentration (100 µM) of ketamine (**Figure 2Aiii** and **B**; MWU; 0 µM: −19±15 spikes, n=6; 100 µM: −49±13 spikes, n=7, p=0.09), compared to control cells that were recorded for the same length of time without ketamine (i.e. 0 µM).

The mean area of the PPs was also substantially reduced (**Figure 2Aii** and **B**), although the difference was not found to be statistically significant within our sample of 7 cells (**Figure 2Ci-ii**; MWU; 0 µM: −26.6±21.7 mV*s, n=6; 100 µM: −80.2±24.6 mV*s, n=7, p=0.45). The effects of ketamine on the plateau and spike number were not readily reversible with wash-out for 5 minutes (**Figure 2B**), perhaps due to the partially lipophilic nature of ketamine (Peltoniemi et al. 2016).

Next, we used prolonged pre-incubation of the slices with different concentrations of ketamine for 2 hours, before inducing PPs with muscarine (**Figure 3**), to test whether the long ketamine incubation would change the effect on the plateaus and spiking. Since ketamine is lipophilic, diffusion from the bath into the slice may be slower than occurs through delivery from capillaries *in vivo*. Thus, Geiger et al (2021) found that in brain slices, ketamine can take up to 40 minutes to fully equilibrate in the tissue. Therefore we decided to use an extended pre-incubation to achieve equilibrium.

We incubated slices in ketamine for 2 hours before transferring them to the recording chamber where the ketamine concentration was maintained, before muscarine was added to elicit a plateau. Plateaus were observed following pre-incubation in 0 µM, 20 µM and 100 µM ketamine (**Figure 3A**), but occurred less often after 100 µM incubation (spiking plateaus in 42% of cells tested) than in control, whereas there was little difference between incubation without ketamine (0 µM: 73% showed plateaus) and with 20 µM ketamine (80% showed plateaus) (**Figure 3B**).

The plateaus in cells incubated in 100 µM ketamine were also significantly smaller than those in cells incubated without (0 µM) and with 20 µM ketamine, with both a smaller area (MWU test; 0 µM: 100.3±15.9 mV*s, n=12; 20 µM: 127.0±24.3 mV*s, n=10; 100 µM: 47.2±11.7 mV*s, n=12; 0 µM vs. 100 µM: p=0.02; 20 µM vs. 100 µM: p=0.02) **(Figure 3Ci**) and fewer spikes (MWU test; 0 µM: 66±12 spikes, n=12; 20 µM: 64±18 spikes, n=10; 100 µM: 17±8.7 spikes, n=12; 0 µM vs. 100 µM: p=0.04; 20 µM vs. 100 µM: p=0.13) (**Figure 3Cii**). The plateaus in cells incubated in 20 µM ketamine were not significantly different in size from those in cells incubated without (0 µM) ketamine, either in area (p=0.28) or spike number (p=0.79). However, the mean values of plateau area and spike frequency were larger after preincubation with 20 µM ketamine than without ketamine (**Figure 3 Ci-ii**). Given our limited cell number (n=12, 10, and 12, for 0, 20, and 100 µM respectively), we do not know whether the surprising, apparent increase in plateau size and spiking was a real, concentration-dependent ketamine effect, or due to chance, including sampling from different subpopulations of L2/3PCs (see discussion).

### Effects of propofol on muscarinic plateaus

We then tested the effects of propofol on the muscarinic plateaus. Wash-in of 3 µM propofol led to an observed slight reduction in the plateau, whilst 10 µM led to a larger reduction (**Figure 4A** and **B**).

Whilst there was no significant difference in the change in the area (MWU test; 0 µM: −18.1±9.5 mV*s, n=10; 3 µM: −25.7±10.0 mV*s, n=8; 10 µM: −57.9±17.8 mV*s, n=10; 0 µM vs. 3 µM: p=0.46; 0 µM vs. 10 µM: p=0.14) following propofol wash-in of either 3 µM or 10 µM compared to 0 µM (**Figure 4Ci**), wash-in of both 3 µM and 10 µM reduced the number of plateau spikes (MWU test; 0 µM: 5.4±9.8 spikes, n=10; 3 µM: −22±8.7 spikes, n=8; 10 µM: −27±12 spikes, n=10; 0 µM vs. 3 µM: p=0.04; 0 µM vs. 10 µM: p=0.048) significantly compared to 0 µM (**Figure 4C**).

As with ketamine we wished to compare the effect on muscarinic plateaus following incubation in propofol, which is of even greater importance as propofol is highly lipophilic, and it has been observed previously that it can take long periods to penetrate into and accumulate in the brain slice preparations similar to those we used (Gredell et al. 2004).

Following incubation in 3 µM propofol plateaus were noticeably smaller (**Figure 5A**), and a smaller proportion of cells exhibited a spiking plateau (spiking plateaus in 44% of cells tested) when compared to cells from slices which had been incubated in standard aCSF (spiking plateaus in 73%; **Figure 5B**). The plateaus in propofol-incubated cells also declined more strongly over time than in control cells (**Figure 5C**). Cells incubated in 3 µM propofol had significantly fewer spikes (MWU test; 0 µM: 66±12 spikes, n=12; 3 µM: 16±5.9 spikes, n=16, p=0.005), but not significantly different post-burst area (MWU test; 0 µM: 100±15.9 mV*s, n=12; 3 µM: 50.1±11.3, n=16, p=0.06) after wash-in of muscarine, compared to those incubated in control aCSF (**Figure 5Di** and **ii**).

## Discussion

In this study, we first confirmed that plateau potentials (PPs), often triggering action potentials, are induced by the metabotropic cholinergic agonist muscarine in layer 2/3 pyramidal cells (L2/3PCs) in slices from the rat prefrontal cortex (PFC), as we found previously (Hagger-Vaughan, Kolnier, and Storm 2023).

The main result of this study, however, is that the general anaesthetics propofol and ketamine attenuate these “muscarinic” PPs in rat PFC L2/3PCs, in a dose-and time-dependent manner. This finding may help explain some of the anaesthetic effects of these drugs *in vivo*.

These results were obtained by testing the anaesthetics before and after bath-applying the arousing agonist muscarine, in order to partly mimic an “awake-like” state in the cortical slices (the “arousal-first approach”). To our knowledge, this approach has not previously been used in *in vitro* experiments with anaesthetics. We used muscarine for this purpose, because muscarinic acetylcholine receptors (mAChR) are known to mediate particularly powerful “arousal” effects in cortical pyramidal cells (Cole and Nicoll 1984; Egorov et al. 2002; Cole and Nicoll 1983).

It is however difficult to clarify how, and to what extent, the effects of anaesthetic drugs in isolated brain slices can be translated into anaesthetic effects of those drugs *in vivo*, including clinical general anaesthesia (GA) in humans. Below we discuss this issue, after first describing the “arousal-first approach” and the induction of PPs by muscarine.

### The “arousal-first approach”: using muscarine to induce an “aroused” state in brain slices before testing anaesthetic drugs

When testing cellular effects of general anaesthetics *in vitro*, one would ideally like to first induce an “aroused”, “awake-like” state in the cells by applying a cocktail of relevant neurochemicals needed for such a state. However, as discussed in the Introduction, it is currently technically impossible to closely mimic, *in vitro*, the complex temporo-spatial patterns of multiple neuromodulators underlying the normal awake state at cortical neurons of the intact brain. Instead, we used a highly simplified, easily controllable “partial arousal-first approach”: adding only a single, powerfully arousing agonist: muscarine. Although not ideal, we think that this approach is considerably more fruitful for testing anaesthetic drugs *in vitro*, than just testing in brain slices or cells that are in a “coma-like” state, due to the profound depletion of nearly all arousing modulation, as have often been done in past *in vitro* studies of anaesthetic drug effects (Hoerbelt and Heifets 2018; Potez and Larkum 2008; Yin et al. 2019; Kitamura et al. 2003; Pittson, Himmel, and MacIver 2004).

Our approach allows testing for effects of anaesthetics on neuronal mechanisms that are likely present in the awake brain *in vivo*, but not found in regular, “comatose” brain slices. In particular, we tested the effects of anaesthetics on the prominent muscarinic, depolarizing PPs, which profoundly alters the input-output function of rat PFC L2/3PCs (Hagger-Vaughan, Kolnier, and Storm 2023). Similarly profound muscarinic effects have also been found in several other cortical pyramidal neuron types (Cole and Nicoll 1983; Egorov et al. 2002). Given the high level of ACh release in the neocortex during conscious brain states *in vivo*, both in wakefulness and dream sleep (Marrosu et al. 1995; Vanini, Lydic, and Baghdoyan 2012; Lee et al. 2005), it seems likely that such PPs in PFC L2/3PCs are a normal feature of the awake, conscious brain, and may contribute strongly to its wake state.

Interestingly, cholinergic/muscarinic “arousal” has previously been used by anaesthesiologists to obtain faster awakening from anaesthesia, by giving cholinesterase inhibitor physostigmine intravenously, thus increasing the ACh level in the brain (Anderson 1988). The effect of this procedure was rather moderate, however, and it is now rarely used. Considering our findings here, the moderate clinical effect of physostigmine under those conditions might perhaps be explained by the remaining anaesthetic (e.g. barbiturates or inhalational agents) still partly suppressing PPs and other cholinergic effects.

### Muscarine reliably induced plateau potentials (PPs) in PFC L2/3PCs

We observed that 10 µM muscarine induced spiking PPs in the majority of PFC L2/3PCs without pre-incubation with an anaesthetic (**Figure 1C**, 73%), thus confirming our previous result (Hagger-Vaughan, Kolnier, and Storm 2023). However, this incidence of eliciting a spiking PP with 10 µM muscarine was lower than the 100% incidence that we found in our previous study. This might be due to the larger number of L2/3PCs cells in this study (n=129) compared to the previous one (n=10; (Hagger-Vaughan, Kolnier, and Storm 2023), as the sample of cells tested here might include a greater diversity of cell phenotypes, i.e. subtypes of L2/3PCs (Yao et al. 2023).

### Comparing relevant drug concentrations and effects *in vitro* vs. *in vivo*. Long pre-incubation revealed effects of lower doses

It is difficult to know which concentrations of anaesthetics to use in slice experiments in order to mimic clinically relevant drug concentrations within the human or animal brain during general anaesthesia. The difficulties stem in part from the complex pharmacokinetics of anaesthetic drugs, including binding to proteins and lipids, as well as diffusion barriers and delays, resulting in large differences between the different compartments (blood vs. brain compartments, water vs. lipid phases, etc.) and between total and effective free drug concentration within each compartment. As it is currently not possible to directly measure anaesthetic drug concentrations at the sites of actions in the human brain, these concentrations have to be inferred from models (Struys et al. 2000). Unfortunately, these issues are often not clearly or explicitly addressed in the available literature. It often seems unclear exactly which concentrations, compartments and conditions authors are referring to. In addition, reported values vary considerably (e.g. Hijazi and Boulieu (2002); Dayton et al. (1983); Ishii-Maruhama et al. (2018); Franks and Lieb (1994). See **Supplement 1** for a more detailed discussion of these issues.

### Effects of ketamine *in vitro* vs. *in vivo*

We saw no significant effect on PPs of relatively low ketamine (20 µM), neither with wash-in nor pre-incubation (**Figures 2** and **3**). In previous *in vitro* experiments with ketamine in rodent CNS slices, a wide range of concentrations have been used, from 30 nM to 10 mM free ketamine (Schnoebel et al. 2005; Yin et al. 2019; Potez and Larkum 2008). Studies in humans have estimated that full unresponsiveness to both verbal command and pain caused by ketamine alone requires at least 3-4 µM total ketamine in the blood (Schüttler et al. 1987), yielding ∼2 µM free ketamine in blood (assuming ∼50% protein binding; Dayton et al. 1983). According to one kind of estimate based on these human data and simplifying assumptions, our *in vitro* concentration of 20 µM free ketamine (**Figures 2** and **3**) may be about 10 times higher than the *in vivo* concentration of free ketamine in the human brain required for full surgical anaesthesia with ketamine alone. However, another estimate, based on *in vivo* data from rodents (Zanos et al. 2016; Levin-Arama et al. 2016; Yang et al. 2018; Cichon et al. 2022), may suggest a free brain concentration of ketamine during full anaesthesia of 20-100 µM, resembling the 20-100 µM free concentrations that we used in the aCSF for our wash-in and pre-incubation experiments (**Figures 2** and **3**). One possibility is that rodents may require higher free brain concentration of ketamine than humans for full anaesthesia (Ribeiro et al. 2014). It should be noted that the IV starting dose to induce general anaesthesia in humans is typically 3-4 mg/kg (Idvall et al. 1979) whereas the IP doses in rodents mentioned above are ∼10-190 mg/kg. The above wide ranges may also reflect that it is very difficult to determine the exact minimal dose of ketamine for full anaesthesia caused by ketamine alone in rodents; hence some reported values may be far too high. In addition, IV injections seem to be much more effective than IP injections of similar ketamine doses in rodents, probably reflecting slower and incomplete absorption into the blood from IP injections. These multiple issues and diverging estimates illustrate the difficulties in choosing appropriate concentrations of anaesthetics for use in slice experiments. See **Supplement 1** for a more detailed discussion.

### Possible relevance of our *in vitro* results with ketamine to anaesthetic and other effects *in vivo*

The estimates based on *in vivo* ketamine anaesthesia in rodents (20-100 µM free ketamine in the brain) suggest that the different effects on PPs) that we found between these concentrations (100 µM ketamine reduced the PPs, whereas 20 µM had little or no effects on the PPs: **Figures 2** and **3**), may be relevant for the *in vivo* anaesthetic effects of ketamine, at least in rodents. Furthermore, this reasoning, based on rodent data, may suggest that the plateau suppression that we saw at higher but not lower ketamine concentrations (**Figures 2** and **3**) may contribute to the suppression of responsiveness combined with altered “inner” consciousness seen at anaesthetic doses in humans (Sarasso et al. 2015).

### Apparent bimodal concentration-dependence of ketamine effects on PPs: possible relevance for consciousness?

We found that the mean values of the PP area and spike frequency were somewhat larger after preincubation with 20 µM ketamine than without ketamine (**Figure 3 Ci-ii**). This might suggest that relatively low ketamine concentrations increased the plateaus whereas higher concentrations reduced the plateaus, thus suggesting a bimodal, concentration-dependent ketamine effect. We do not know whether this surprising pattern is real or due to chance, or might reflect different subpopulations of L2/3PCs that we could not distinguish due to small sample sizes (van Aerde and Feldmeyer 2015). However, if there is really such a bimodal concentration-dependence of the ketamine effect on the PPs, it seems to fit well with, and may help explain, certain observations of seemingly bimodal, concentration-dependent effects of ketamine *in vivo* both in humans (Farnes et al. 2020) and in rodents (Arena et al. 2022).

The apparent enhancing effect on PPs of relatively low ketamine (20 µM) in both wash-in and pre-incubation experiments (**Figures 2** and **3**), may suggest that low ketamine doses may exert their potential antidepressant effects (Mion and Villevieille 2013) by enhancing PPs and persistent firing. This possibility may be supported by evidence suggesting that sub-anaesthetic, antidepressant ketamine doses cause a hyperglutamatergic state in the PFC, in contrast to anaesthetic doses which decrease PFC glutamate levels (Moghaddam et al. 1997). This again suggests a bimodal concentration-dependence, which might be related to an enhanced PP-driven firing and glutamate release induced by low ketamine (**Figure 3 Ci-ii**).

### Effects of propofol *in vitro* vs. *in vivo*

The total blood plasma concentrations of propofol during surgical anaesthesia in humans have been estimated to ∼3-27 mg/ml (C. Smith et al. 1994; Braathen et al. 2022), which converts to ∼16-153 µM. However, this range reflects large differences between “unresponsiveness” to speech (required 5.4 µg/ml propofol) vs. to pain (required 27 µg/ml) induced by propofol alone. Since propofol is ∼95-97% bound to plasma proteins (Cockshott et al. 1992), the 3 µM propofol used in our ACSF for slice experiments (**Figures 4** and **5**) may be close to the concentration range relevant in humans for deep general anaesthesia and analgesia with propofol only. However, this dosage is almost never used for clinical surgery, as propofol is normally used at 3-4 µg/ml total in plasma (∼0.5 µM free propofol) for verbal unconsciousness, supplemented with another drug for analgesic unresponsiveness. In general, available evidence suggests that *in vitro* experiments often require higher drug concentrations than *in vivo*, but the mechanisms are complex and often poorly understood (Leist et al. 2017; Hengstler et al. 2020).

### Long pre-incubation of propofol was needed to reveal effects of lower doses

A major difference between intravenous administration of ketamine or propofol *in vivo* and our bath-application of these drugs *in vitro*, is that brain slices of course lack the *in vivo* blood supply through a dense capillary net, which brings drugs rapidly to within a few micrometres from each brain cell, requiring minimal diffusion time. This difference proved to be particularly important for propofol and probably explains its far slower equilibration in slices.

In line with this reasoning, our experiments using 2 hours of pre-incubation with relatively low concentrations of propofol (3 µM) or ketamine (20 µM) showed considerably stronger drug effects than our wash-in experiments applying the same drug concentrations for 5-25 minutes. This difference was especially striking for propofol (compare **Figures 2** and **3** for ketamine, and **Figures 4** and **5** for propofol). This strongly suggests that our wash-in applications did not reveal the full effect of the applied drug concentrations, presumably because of delayed equilibration between the medium and the slice. This interpretation is strongly supported by studies of diffusion of ketamine and propofol (Gredell et al. 2004; Geiger et al. 2021). Therefore, our discussion and conclusions are primarily based on our 2 h pre-incubation results.

### Effects of ketamine and propofol on PPs vs. states of consciousness

How may the similarity in results between high concentrations of ketamine and propofol (100 µM and 10 µM respectively, **Figures 2** and **4**) be related to their dramatically different states of unconsciousness? The similarity between the effects of ketamine and propofol that we observed suggests that PPs may have a greater role in supporting external awareness and responsiveness than on conscious experience itself, as reportable “inner” consciousness in the form of vivid dreaming is preserved in ketamine but less so in propofol anaesthesia, whilst responsiveness are abolished in both (Sarasso et al. 2015; Nilsen et al. 2022).

## Summary and conclusions

We used whole-cell patch clamp recordings to compare the effects of two different general anaesthetics, ketamine and propofol, in L2/3 pyramidal cells (PCs) in prefrontal rat cortical brain slices. We pre-treated the slices by first inducing a partially “aroused” state by adding muscarine, which caused depolarizing plateau potentials (PPs) with spiking.

After a long (2 hour) pre-incubation with 20 µM ketamine or 3 µM propofol, assumed to be close to equipotent doses in rats, the muscarine-induced PPs were altered in different ways: 3 µM propofol reduced the mean PPs area (non-significant) and significantly reduced the spiking, whereas 20 µM ketamine was followed by an apparent (non-significant) increase in the PPs.

The apparent differential effects on PPs of our lower concentrations of ketamine (20 µM) or propofol (3 µM) may be related to their contrasting clinical effects: ketamine causing atypical anaesthesia (unresponsiveness) with desynchronized cortical activity and often vivid, “psychedelic” dreaming, whereas propofol causes slow-wave activity with unresponsiveness and less dreaming.

We also tested a briefer wash-in (5-20 min) of ketamine or propofol. Higher concentrations of ketamine (100 µM) or propofol (10 µM), which are known to cause deeper anaesthesia *in vivo*, both suppressed PPs and spiking, suggesting a possible relation between PPs, desynchronized cortical activity, and conscious experience. Propofol (3 µM) but not ketamine (20 µM) significantly reduced the spike frequency, and we found no significant changes in mean PPs, perhaps reflecting insufficient equilibration by diffusion into the slices.

Choosing concentrations and timing of anaesthetics for *in vitro* slice experiments is a complex issue, and much remains to be clarified as to relevance for *in vivo* human anaesthesia. Based on simplifying, uncertain assumptions, the concentrations we used *in vitro* appear to be somewhat higher than those normally used in clinical human anaesthesia. More experiments are needed to test our tentative conclusions, including the apparent enhancement of PPs by relatively low ketamine.

## Data availability statement

The data that support the findings of this study are available from the corresponding author, JFS, upon request.

## Ethics statement

All animal procedures were approved by the responsible veterinarian of the institute, in accordance with the statute regulating animal experimentation (Norwegian Ministry of Agriculture, 1996). Ethical review and approval by an ethics committee was not required for this animal study, because such a specific ethical approval is not required for individual studies utilising only *ex vivo* experiments.

## Author contributions

JFS suggested the “arousal first” approach and the ketamine experiments to ASF for a MSRP project, and the propofol experiments for NHV’s PhD project. ASF, DK and NHV conducted the experiments and analysed the data. JFS supervised the study and provided the funding. JR provided information about and discussed clinically relevant concentrations of anaesthetics in humans. All helped writing and revising the manuscript.

## Acknowledgements

We would like to thank our colleague Dr. Christoph Hönigsperger for comments, and Cecile Oksvold for technical assistance.

## References

Aerde, Karlijn I. van, and Dirk Feldmeyer. 2015. “Morphological and Physiological Characterization of Pyramidal Neuron Subtypes in Rat Medial Prefrontal Cortex.” Cerebral Cortex 25 (3): 788–805.

Anderson, J. A. 1988. “Reversal Agents in Sedation and Anesthesia: A Review.” Anesthesia Progress 35 (2): 43–47.

Arena, A., B. E. Juel, R. Comolatti, S. Thon, and J. F. Storm. 2022. “Capacity for Consciousness under Ketamine Anaesthesia Is Selectively Associated with Activity in Posteromedial Cortex in Rats.” Neuroscience of Consciousness 2022 (1): niac004.

Aru, Jaan, Francesca Siclari, William A. Phillips, and Johan F. Storm. 2020. “Apical Drive-A Cellular Mechanism of Dreaming?” Neuroscience and Biobehavioral Reviews 119 (December): 440–55.

Aru, Jaan, Mototaka Suzuki, and Matthew E. Larkum. 2021. “Cellular Mechanisms of Conscious Processing: (Trends in Cognitive Sciences, 24, 814-825; 2020).” Trends in Cognitive Sciences 25 (12): 1096.

Asaad, Wael F., Gregor Rainer, and Earl K. Miller. 1998. “Neural Activity in the Primate Prefrontal Cortex during Associative Learning.” Neuron 21 (6): 1399–1407.

Bahnasi, Y. M., H. M. Wright, C. J. Milligan, A. M. Dedman, F. Zeng, P. M. Hopkins, A. N. Bateson, and D. J. Beech. 2008. “Modulation of TRPC5 Cation Channels by Halothane, Chloroform and Propofol.” British Journal of Pharmacology 153 (7): 1505–12.

Bliss, T. V. P., and G. L. Collingridge. 1993. “A Synaptic Model of Memory: Long-Term Potentiation in the Hippocampus.” Nature 361 (6407): 31–39.

Boly, Melanie, Marcello Massimini, Naotsugu Tsuchiya, Bradley R. Postle, Christof Koch, and Giulio Tononi. 2017. “Are the Neural Correlates of Consciousness in the Front or in the Back of the Cerebral Cortex? Clinical and Neuroimaging Evidence.” The Journal of Neuroscience: The Official Journal of the Society for Neuroscience 37 (40): 9603–13.

Braathen, Martin R., Ivan Rimstad, Terje Dybvik, Ståle Nygård, and Johan Raeder. 2022. “Online Exhaled Propofol Monitoring in Normal-Weight and Obese Surgical Patients.” Acta Anaesthesiologica Scandinavica 66 (5): 598–605.

Brown, Richard, Hakwan Lau, and Joseph E. LeDoux. 2019. “Understanding the Higher-Order Approach to Consciousness.” Trends in Cognitive Sciences 23 (9): 754–68.

Burgos-Robles, Anthony, Eyal Y. Kimchi, Ehsan M. Izadmehr, Mary Jane Porzenheim, William A. Ramos-Guasp, Edward H. Nieh, Ada C. Felix-Ortiz, et al. 2017. “Amygdala Inputs to Prefrontal Cortex Guide Behavior amid Conflicting Cues of Reward and Punishment.” Nature Neuroscience 20 (6): 824–35.

Casali, Adenauer G., Olivia Gosseries, Mario Rosanova, Mélanie Boly, Simone Sarasso, Karina R. Casali, Silvia Casarotto, et al. 2013. “A Theoretically Based Index of Consciousness Independent of Sensory Processing and Behavior.” Science Translational Medicine 5 (198): 198ra105.

Casey, Cameron P., Sean Tanabe, Zahra Farahbakhsh, Margaret Parker, Amber Bo, Marissa White, Tyler Ballweg, et al. 2022. “Distinct EEG Signatures Differentiate Unconsciousness and Disconnection during Anaesthesia and Sleep.” British Journal of Anaesthesia 128 (6): 1006–18.

Chen, Xiangdong, Shaofang Shu, and Douglas A. Bayliss. 2009. “HCN1 Channel Subunits Are a Molecular Substrate for Hypnotic Actions of Ketamine.” The Journal of Neuroscience: The Official Journal of the Society for Neuroscience 29 (3): 600–609.

Cichon, Joseph, Andrzej Z. Wasilczuk, Loren L. Looger, Diego Contreras, Max B. Kelz, and Alex Proekt. 2022. “Ketamine Triggers a Switch in Excitatory Neuronal Activity across Neocortex.” Nature Neuroscience 26 (1): 39–52.

Clark, Kelsey, Ryan Fox Squire, Yaser Merrikhi, and Behrad Noudoost. 2015. “Visual Attention: Linking Prefrontal Sources to Neuronal and Behavioral Correlates.” Progress in Neurobiology 132 (September): 59–80.

Cockshott, I. D., E. J. Douglas, G. F. Plummer, and P. J. Simons. 1992. “The Pharmacokinetics of Propofol in Laboratory Animals.” Xenobiotica; the Fate of Foreign Compounds in Biological Systems 22 (3): 369–75.

Cole, A. E., and R. A. Nicoll. 1983. “Acetylcholine Mediates a Slow Synaptic Potential in Hippocampal Pyramidal Cells.” Science 221 (4617): 1299–1301.

Cole, A. E., and R. A. Nicoll. 1984. “Characterization of a Slow Cholinergic Post-Synaptic Potential Recorded in Vitro from Rat Hippocampal Pyramidal Cells.” The Journal of Physiology 352 (July). 10.1113/jphysiol.1984.sp015285.

Collier, B. B. 1972. “Ketamine and the Conscious Mind.” Anaesthesia 27 (2): 120–34.

Colombo, Michele Angelo, Martino Napolitani, Melanie Boly, Olivia Gosseries, Silvia Casarotto, Mario Rosanova, Jean-Francois Brichant, et al. 2019. “The Spectral Exponent of the Resting EEG Indexes the Presence of Consciousness during Unresponsiveness Induced by Propofol, Xenon, and Ketamine.” NeuroImage 189 (April): 631–44.

D’Andola, Mattia, Beatriz Rebollo, Adenauer G. Casali, Julia F. Weinert, Andrea Pigorini, Rosa Villa, Marcello Massimini, and Maria V. Sanchez-Vives. 2018. “Bistability, Causality, and Complexity in Cortical Networks: An In Vitro Perturbational Study.” Cerebral Cortex 28 (7): 2233–42.

Dasilva, Miguel, Alessandra Camassa, Alvaro Navarro-Guzman, Antonio Pazienti, Lorena Perez-Mendez, Gorka Zamora-López, Maurizio Mattia, and Maria V. Sanchez-Vives. 2021. “Modulation of Cortical Slow Oscillations and Complexity across Anesthesia Levels.” NeuroImage 224 (January): 117415.

Dayton, P. G., R. L. Stiller, D. R. Cook, and J. M. Perel. 1983. “The Binding of Ketamine to Plasma Proteins: Emphasis on Human Plasma.” European Journal of Clinical Pharmacology 24 (6): 825–31.

Dehaene, S., M. Kerszberg, and J. P. Changeux. 1998. “A Neuronal Model of a Global Workspace in Effortful Cognitive Tasks.” Proceedings of the National Academy of Sciences of the United States of America 95 (24): 14529–34.

Dingledine, Raymond. 1984. Brain Slices. Springer US.

Edlow, Brian L., Jan Claassen, Nicholas D. Schiff, and David M. Greer. 2020. “Recovery from Disorders of Consciousness: Mechanisms, Prognosis and Emerging Therapies.” Nature Reviews. Neurology 17 (3): 135–56.

Edwards, Zachary, and Charles V. Preuss. 2023. “GABA Receptor Positive Allosteric Modulators.” In StatPearls. Treasure Island (FL): StatPearls Publishing.

Egorov, Alexei V., Bassam N. Hamam, Erik Fransén, Michael E. Hasselmo, and Angel A. Alonso. 2002. “Graded Persistent Activity in Entorhinal Cortex Neurons.” Nature 420 (6912): 173–78.

Farnes, Nadine, Bjørn E. Juel, André S. Nilsen, Luis G. Romundstad, and Johan F. Storm. 2020. “Increased Signal Diversity/complexity of Spontaneous EEG, but Not Evoked EEG Responses, in Ketamine-Induced Psychedelic State in Humans.” PloS One 15 (11): e0242056.

Fassl, Jens, Kane M. High, Edward R. Stephenson, Viktor Yarotskyy, and Keith S. Elmslie. 2011. “The Intravenous Anesthetic Propofol Inhibits Human L-Type Calcium Channels by Enhancing Voltage-Dependent Inactivation.” Journal of Clinical Pharmacology 51 (5): 719–30.

Franks, N. P. 2008. “General Anaesthesia: From Molecular Targets to Neuronal Pathways of Sleep and Arousal.” Nature Reviews. Neuroscience 9 (5): 370–86.

Franks, N. P., and W. R. Lieb. 1994. “Molecular and Cellular Mechanisms of General Anaesthesia.” Nature 367 (6464): 607–14.

Fraser, D. D., and B. A. MacVicar. 1996. “Cholinergic-Dependent Plateau Potential in Hippocampal CA1 Pyramidal Neurons.” The Journal of Neuroscience: The Official Journal of the Society for Neuroscience 16 (13): 4113–28.

Gambino, Frédéric, Stéphane Pagès, Vassilis Kehayas, Daniela Baptista, Roberta Tatti, Alan Carleton, and Anthony Holtmaat. 2014. “Sensory-Evoked LTP Driven by Dendritic Plateau Potentials in Vivo.” Nature 515 (7525): 116–19.

Garcia-Rill, Edgar, Nebojsa Kezunovic, James Hyde, Christen Simon, Paige Beck, and Francisco J. Urbano. 2013. “Coherence and Frequency in the Reticular Activating System (RAS).” Sleep Medicine Reviews 17 (3): 227–38.

Geiger, Zachary, Brett VanVeller, Zarin Lopez, Abdel K. Harrata, Kathryn Battani, Lauren Wegman-Points, and Li-Lian Yuan. 2021. “Determination of Diffusion Kinetics of Ketamine in Brain Tissue: Implications for in Vitro Mechanistic Studies of Drug Actions.” Frontiers in Neuroscience. 10.3389/fnins.2021.678978.

Goldman-Rakic, P. S. 1988. “Topography of Cognition: Parallel Distributed Networks in Primate Association Cortex.” Annual Review of Neuroscience 11: 137–56.

Grace, R. F. 2003. “The Effect of Variable-Dose Diazepam on Dreaming and Emergence Phenomena in 400 Cases of Ketamine-Fentanyl Anaesthesia.” Anaesthesia 58 (9): 904–10.

Gredell, J. A., P. A. Turnquist, M. B. Maciver, and R. A. Pearce. 2004. “Determination of Diffusion and Partition Coefficients of Propofol in Rat Brain Tissue: Implications for Studies of Drug Action in Vitro.” British Journal of Anaesthesia 93 (6): 810–17.

Hagger-Vaughan, Nicholas, Daniel Kolnier, and Johan F. Storm. 2023. “Non-Apical Plateau Potentials and Persistent Firing Induced by Metabotropic Cholinergic Modulation in Layer 2/3 Pyramidal Cells in the Rat Prefrontal Cortex.” bioRxiv. 10.1101/2023.11.02.565356.

Hagger-Vaughan, Nicholas, and Johan F. Storm. 2019. “Synergy of Glutamatergic and Cholinergic Modulation Induces Plateau Potentials in Hippocampal OLM Interneurons.” Frontiers in Cellular Neuroscience 13 (November): 508.

Hengstler, Jan G., Anna-Karin Sjögren, Daniele Zink, and Jorrit J. Hornberg. 2020. “In Vitro Prediction of Organ Toxicity: The Challenges of Scaling and Secondary Mechanisms of Toxicity.” Archives of Toxicology 94 (2): 353–56.

Hijazi, Youssef, and Roselyne Boulieu. 2002. “Protein Binding of Ketamine and Its Active Metabolites to Human Serum.” European Journal of Clinical Pharmacology 58 (1): 37–40.

Hobson, J. Allan. 2009. “REM Sleep and Dreaming: Towards a Theory of Protoconsciousness.” Nature Reviews. Neuroscience 10 (11): 803–13.

Hoerbelt, Paul, and Boris D. Heifets. 2018. “Native System and Cultured Cell Electrophysiology for Investigating Anesthetic Mechanisms.” Methods in Enzymology 602 (March): 301–38.

Hofschneider, Mark. 2018. “Discovery and Development of Propofol, a Widely Used Anesthetic.” Lasker Foundation. September 10, 2018. https://laskerfoundation.org/winners/discovery-and-development-of-propofol-a-widely-used-anesthetic/.

Idvall, J., I. Ahlgren, K. R. Aronsen, and P. Stenberg. 1979. “Ketamine Infusions: Pharmacokinetics and Clinical Effects.” British Journal of Anaesthesia 51 (12): 1167–73.

Ishii-Maruhama, Minako, Hitoshi Higuchi, Mai Nakanou, Yuka Honda-Wakasugi, Akiko Yabuki-Kawase, Shigeru Maeda, and Takuya Miyawaki. 2018. “In Vitro Changes in the Proportion of Protein-Unbound-Free Propofol Induced by Valproate.” Journal of Anesthesia 32 (5): 688–93.

Kitamura, Akira, William Marszalec, Jay Z. Yeh, and Toshio Narahashi. 2003. “Effects of Halothane and Propofol on Excitatory and Inhibitory Synaptic Transmission in Rat Cortical Neurons.” The Journal of Pharmacology and Experimental Therapeutics 304 (1): 162–71.

Kurdi, Madhuri S., Kaushic A. Theerth, and Radhika S. Deva. 2014. “Ketamine: Current Applications in Anesthesia, Pain, and Critical Care.” Anesthesia, Essays and Researches 8 (3): 283–90.

Lee, Maan Gee, Oum K. Hassani, Angel Alonso, and Barbara E. Jones. 2005. “Cholinergic Basal Forebrain Neurons Burst with Theta during Waking and Paradoxical Sleep.” The Journal of Neuroscience: The Official Journal of the Society for Neuroscience 25 (17): 4365–69.

Leist, Marcel, Ahmed Ghallab, Rabea Graepel, Rosemarie Marchan, Reham Hassan, Susanne Hougaard Bennekou, Alice Limonciel, et al. 2017. “Adverse Outcome Pathways: Opportunities, Limitations and Open Questions.” Archives of Toxicology 91 (11): 3477–3505.

Levin-Arama, Maya, Lital Abraham, Trevor Waner, Alon Harmelin, David M. Steinberg, Tal Lahav, and Mickey Harlev. 2016. “Subcutaneous Compared with Intraperitoneal KetamineXylazine for Anesthesia of Mice.” Journal of the American Association for Laboratory Animal Science: JAALAS 55 (6): 794–800.

Lo, Fu-Sun, and Reha S. Erzurumlu. 2002. “L-Type Calcium Channel-Mediated Plateau Potentials in Barrelette Cells during Structural Plasticity.” Journal of Neurophysiology 88 (2): 794–801.

Lydic, Ralph, and Helen A. Baghdoyan. 2002. “Ketamine and MK-801 Decrease Acetylcholine Release in the Pontine Reticular Formation, Slow Breathing, and Disrupt Sleep.” Sleep 25 (6): 617–22.

Marrosu, Francesco, Chiara Portas, Maria Stefania Mascia, Maria Antonietta Casu, Mauro Fà, Marcello Giagheddu, Assunta Imperato, and Gian Luigi Gessa. 1995. “Microdialysis Measurement of Cortical and Hippocampal Acetylcholine Release during Sleep-Wake Cycle in Freely Moving Cats.” Brain Research 671 (2): 329–32.

Mashour, George A., Pieter Roelfsema, Jean-Pierre Changeux, and Stanislas Dehaene. 2020. “Conscious Processing and the Global Neuronal Workspace Hypothesis.” Neuron 105 (5): 776–98.

Maurer, F., D. J. Lorenz, G. Pielsticker, T. Volk, D. I. Sessler, J. I. Baumbach, and S. Kreuer. 2017. “Adherence of Volatile Propofol to Various Types of Plastic Tubing.” Journal of Breath Research 11 (1): 016009.

McCormick, D. A., and D. A. Prince. 1985. “Two Types of Muscarinic Response to Acetylcholine in Mammalian Cortical Neurons.” Proceedings of the National Academy of Sciences of the United States of America 82 (18): 6344–48.

Mion, Georges, and Thierry Villevieille. 2013. “Ketamine Pharmacology: An Update (Pharmacodynamics and Molecular Aspects, Recent Findings).” CNS Neuroscience & Therapeutics 19 (6): 370–80.

Moghaddam, B., B. Adams, A. Verma, and D. Daly. 1997. “Activation of Glutamatergic Neurotransmission by Ketamine: A Novel Step in the Pathway from NMDA Receptor Blockade to Dopaminergic and Cognitive Disruptions Associated with the Prefrontal Cortex.” The Journal of Neuroscience: The Official Journal of the Society for Neuroscience 17 (8): 2921–27.

Moliner, Rafael, Mykhailo Girych, Cecilia A. Brunello, Vera Kovaleva, Caroline Biojone, Giray Enkavi, Lina Antenucci, et al. 2023. “Psychedelics Promote Plasticity by Directly Binding to BDNF Receptor TrkB.” Nature Neuroscience 26 (6): 1032–41.

Moore, Tirin, and Katherine M. Armstrong. 2003. “Selective Gating of Visual Signals by Microstimulation of Frontal Cortex.” Nature 421 (6921): 370–73.

Moruzzi, G., and H. W. Magoun. 1949. “Brain Stem Reticular Formation and Activation of the EEG.” Electroencephalography and Clinical Neurophysiology 1 (1-4): 455–73.

Nilsen, André S., Alessandro Arena, and Johan F. Storm. 2023. “Effect of Anesthesia in Rats on Measures of Complexity, Differentiation, and Integrated Information.” PsyArXiv. 10.31234/osf.io/untrm.

Nilsen, André S., Bjørn E. Juel, Benjamin Thürer, Arnfinn Aamodt, and Johan F. Storm. 2022. “Are We Really Unconscious in ‘unconscious’ States? Common Assumptions Revisited.” Frontiers in Human Neuroscience 16 (October): 987051.

Nordström, O., A. M. Engström, S. Persson, and R. Sandin. 1997. “Incidence of Awareness in Total I.v. Anaesthesia Based on Propofol, Alfentanil and Neuromuscular Blockade.” Acta Anaesthesiologica Scandinavica 41 (8): 978–84.

Nuñez, Angel, Soledad Domínguez, Washington Buño, and David Fernández de Sevilla. 2012. “Cholinergic-Mediated Response Enhancement in Barrel Cortex Layer V Pyramidal Neurons.” Journal of Neurophysiology 108 (6): 1656–68.

Oikonomou, Katerina D., Mandakini B. Singh, Enas V. Sterjanaj, and Srdjan D. Antic. 2014. “Spiny Neurons of Amygdala, Striatum, and Cortex Use Dendritic Plateau Potentials to Detect Network UP States.” Frontiers in Cellular Neuroscience 8 (September): 292.

Paus, Tomáš. 2000. “Functional Anatomy of Arousal and Attention Systems in the Human Brain” 126 (January): 65–77.

Peltoniemi, Marko A., Nora M. Hagelberg, Klaus T. Olkkola, and Teijo I. Saari. 2016. “Ketamine: A Review of Clinical Pharmacokinetics and Pharmacodynamics in Anesthesia and Pain Therapy.” Clinical Pharmacokinetics 55 (9): 1059–77.

Pittson, Sky, Allison M. Himmel, and M. Bruce MacIver. 2004. “Multiple Synaptic and Membrane Sites of Anesthetic Action in the CA1 Region of Rat Hippocampal Slices.” BMC Neuroscience 5 (December): 52.

Posner, Jerome B., Clifford B. Saper, Nicholas Schiff, and Fred Plum. 2007. Plum and Posner’s Diagnosis of Stupor and Coma. Oxford University Press.

Potez, Sarah, and Matthew E. Larkum. 2008. “Effect of Common Anesthetics on Dendritic Properties in Layer 5 Neocortical Pyramidal Neurons.” Journal of Neurophysiology 99 (3): 1394–1407.

Ribeiro, Patrícia O., Angelo R. Tomé, Henrique B. Silva, Rodrigo A. Cunha, and Luís M. Antunes. 2014. “Clinically Relevant Concentrations of Ketamine Mainly Affect Long-Term Potentiation rather than Basal Excitatory Synaptic Transmission and Do Not Change Paired-Pulse Facilitation in Mouse Hippocampal Slices.” Brain Research 1560 (April): 10–17.

Sanchez-Vives, Maria V., Marcello Massimini, and Maurizio Mattia. 2017. “Shaping the Default Activity Pattern of the Cortical Network.” Neuron 94 (5): 993–1001.

Sanders, Robert D., Giulio Tononi, Steven Laureys, and Jamie W. Sleigh. 2012. “Unresponsiveness ≠ Unconsciousness.” Anesthesiology 116 (4): 946–59.

Sarasso, Simone, Melanie Boly, Martino Napolitani, Olivia Gosseries, Vanessa Charland-Verville, Silvia Casarotto, Mario Rosanova, et al. 2015. “Consciousness and Complexity during Unresponsiveness Induced by Propofol, Xenon, and Ketamine.” Current Biology: CB 25 (23): 3099–3105.

Schnoebel, Rose, Matthias Wolff, Saskia C. Peters, Michael E. Bräu, Andreas Scholz, Gunter Hempelmann, Horst Olschewski, and Andrea Olschewski. 2005. “Ketamine Impairs Excitability in Superficial Dorsal Horn Neurones by Blocking Sodium and Voltage-Gated Potassium Currents.” British Journal of Pharmacology 146 (6): 826–33.

Smith, C., A. I. McEwan, R. Jhaveri, M. Wilkinson, D. Goodman, L. R. Smith, A. T. Canada, and P. S. Glass. 1994. “The Interaction of Fentanyl on the Cp50 of Propofol for Loss of Consciousness and Skin Incision.” Anesthesiology 81 (4): 820–28; discussion 26A.

Smith, Guerin, Jason R. D’Cruz, Bryan Rondeau, and Julie Goldman. 2023. “General Anesthesia for Surgeons.” In StatPearls. Treasure Island (FL): StatPearls Publishing.

Steriade, Mircea. 2001. The Intact and Sliced Brain. MIT Press.

Steriade, I. Timofeev, and F. Grenier. 2001. “Natural Waking and Sleep States: A View From Inside Neocortical Neurons.” *Journal of Neurophysiology*, May. 10.1152/jn.2001.85.5.1969.

Struys, M. M., T. De Smet, B. Depoorter, L. F. Versichelen, E. P. Mortier, F. J. Dumortier, S. L. Shafer, and G. Rolly. 2000. “Comparison of Plasma Compartment versus Two Methods for Effect Compartment--Controlled Target-Controlled Infusion for Propofol.” Anesthesiology 92 (2): 399–406.

Timofeev, I., F. Grenier, M. Bazhenov, T. J. Sejnowski, and M. Steriade. 2000. “Origin of Slow Cortical Oscillations in Deafferented Cortical Slabs.” Cerebral Cortex 10 (12): 1185–99.

Trapani, G., C. Altomare, G. Liso, E. Sanna, and G. Biggio. 2000. “Propofol in Anesthesia. Mechanism of Action, Structure-Activity Relationships, and Drug Delivery.” Current Medicinal Chemistry 7 (2): 249–71.

Vanini, Giancarlo, Ralph Lydic, and Helen A. Baghdoyan. 2012. “GABA-to-ACh Ratio in Basal Forebrain and Cerebral Cortex Varies Significantly during Sleep.” Sleep 35 (10): 1325–34.

Wang, Xiao-Jing. 2001. “Synaptic Reverberation Underlying Mnemonic Persistent Activity.” Trends in Neurosciences 24 (8): 455–63.

Watson, Christopher J., Helen A. Baghdoyan, and Ralph Lydic. 2010. “Neuropharmacology of Sleep and Wakefulness.” Sleep Medicine Clinics 5 (4): 513–28.

White, P. F., W. L. Way, and A. J. Trevor. 1982. “Ketamine--Its Pharmacology and Therapeutic Uses.” Anesthesiology 56 (2): 119–36.

Williams, Stephen R., and Lee N. Fletcher. 2019. “A Dendritic Substrate for the Cholinergic Control of Neocortical Output Neurons.” Neuron 101 (3): 486–99.e4.

Yamakage, Michiaki, Carol A. Hirshman, and Thomas L. Croxton. 1995. “Inhibitory Effects of Thiopental, Ketamine, and Propofol on Voltage-Dependent Calcium Sup 2+ Channels in Porcine Tracheal Smooth Muscle Cells.” Anesthesiology 83 (6): 1274–82.

Yang, Yan, Yihui Cui, Kangning Sang, Yiyan Dong, Zheyi Ni, Shuangshuang Ma, and Hailan Hu. 2018. “Ketamine Blocks Bursting in the Lateral Habenula to Rapidly Relieve Depression.” Nature 554 (7692): 317–22.

Yao, Zizhen, Cindy T. J. van Velthoven, Michael Kunst, Meng Zhang, Delissa McMillen, Changkyu Lee, Won Jung, et al. 2023. “A High-Resolution Transcriptomic and Spatial Atlas of Cell Types in the Whole Mouse Brain.” Nature 624 (7991): 317–32.

Ying, Shui-Wang, Syed Y. Abbas, Neil L. Harrison, and Peter A. Goldstein. 2006. “Propofol Block of I(h) Contributes to the Suppression of Neuronal Excitability and Rhythmic Burst Firing in Thalamocortical Neurons.” The European Journal of Neuroscience 23 (2): 465–80.

Ying, Shui-Wang, and Peter A. Goldstein. 2005. “Propofol-Block of SK Channels in Reticular Thalamic Neurons Enhances GABAergic Inhibition in Relay Neurons.” Journal of Neurophysiology 93 (4): 1935–48.

Yin, Jianyin, Bao Fu, Yuan Wang, and Tian Yu. 2019. “Effects of Ketamine on Voltage-Gated Sodium Channels in the Barrel Cortex and the Ventral Posteromedial Nucleus Slices of Rats.” Neuroreport 30 (17): 1197–1204.

Zanos, Panos, Ruin Moaddel, Patrick J. Morris, Polymnia Georgiou, Jonathan Fischell, Greg I. Elmer, Manickavasagom Alkondon, et al. 2016. “NMDAR Inhibition-Independent Antidepressant Actions of Ketamine Metabolites.” Nature 533 (7604): 481–86.

