## Supplemental text 1 for "Effects of ketamine and propofol on muscarinic plateau potentials in rat neocortical pyramidal cells"

### Supplement 1

In this supplement, we provide a more full version of the part of the discussion that concerns our comparison of relevant drug concentrations and their effects *in vitro* vs. *in vivo*, with more extensive and detailed arguments and references. As these issues are important for the clinical relevance of our *in vitro* results, and are also quite complex, they require more space and details than can reasonably be used within the general Discussion.

#### General considerations

For ketamine, the dose needed in humans for unresponsiveness to pain is approximately similar to the dose needed for unresponsiveness to verbal command or mild stimuli (Idvall et al. 1979). For propofol, this is very different as the dose needed for no response to pain is 4-5 times higher than the dose needed for profound sleep or unresponsiveness to verbal command (Smith et al. 1994). Thus, for clinical surgical anaesthesia in humans, propofol is used in a moderate dose together with strong analgesic agents (e.g. opioids) whereas ketamine may be used alone.

The use of isolated brain slices to study the effects of general anaesthetics faces other problems as well, in addition to the “comatose slice” issue. It is difficult to know which concentrations of anaesthetics to use in slice experiments in order to mimic clinically relevant drug concentrations within the human or animal brain during general anaesthesia. A variety of factors are at play, including *in vivo* vs. *in vitro* conditions; different compartments and phases: blood, CSF, aCSF, slices, water/lipid; varying binding to proteins, lipids, and membranes; diffusion barriers and time-dependence; differences between blood-perfused living organs and passive drug diffusion into thick slices; species differences (humans vs. rodents); different or poorly defined criteria and end points for “unconscious” and “unresponsive” states; possible degradation and binding to plastics and other materials in vials and tubes, etc.

##### Pharmacokinetics

The difficulties stem in part from the complex pharmacokinetics of anaesthetic drugs, including binding to proteins and lipids, as well as diffusion barriers and delays, resulting in large differences both between the different compartments (blood vs. brain compartments, water vs. lipid phases, etc.) and between total and effective free drug concentration within each compartment. As it is currently not possible to directly measure anaesthetic drug concentrations at the sites of actions in the human brain, these concentrations have to be inferred from models (Struys et al. 2000). *In vivo*, the half-time needed for equilibration between blood and brain cells has been estimated to be about 2 min. for propofol, and 3-4 min. for ketamine (Struys et al. 2000; Schüttler et al. 1987). Thus, 2 minutes after ketamine reaches the brain capillaries, the free ketamine concentration around the neurons should be about 67% of the free ketamine concentration in blood, and nearly at full equilibrium after ~10 minutes (Struys et al. 2000; Schüttler et al. 1987).

##### Lack of clarity in available literature

Unfortunately, these issues are often not clearly, explicitly addressed in the available literature. It often seems unclear exactly which concentrations, compartments and conditions authors are referring to. In addition, the reported values for anaesthetic drug concentrations, their protein binding etc. vary considerably (e-g. 60-64% protein binding of ketamine in human serum according to Hijazi and Boulieu (2002) vs. about 20-30% in human plasma according to Dayton et al. (1983). The protein-binding of propofol in blood is ~95-96% according to Ishii-Maruhama et al. (2018), but ~97-98 % according to Franks and Lieb (1994). We explicitly discuss some of the most relevant issues below, and will review these problems in a separate paper.

Clinical anaesthesia, procedures and considerations. For clinical anaesthesia, propofol and ketamine are administered by i.v. injection of doses that are normally calculated in mg of drug per kg of body mass, and given as bolus injections and/or as steady infusions at appropriate rates, which are titrated as needed based on the observed level of anaesthesia (Pai and Heining 2007; Scheepstra et al. 1989). The dose required to produce anaesthesia can vary considerably between individuals, and also varies with sex and age (Searle and Hopkins 2009; Braithwaite et al. 2023; Scheepstra et al. 1989). Once these drugs have entered the blood-stream, they partly bind to plasma proteins, and the free fraction then passes the blood-brain barrier, diffuse through the extracellular space and potentially partition into or pass through neuronal membranes before exerting their effects on neuronal molecules and properties (Dayton et al. 1983). All of these factors complicate estimation of the relevant local concentrations in the brain during anaesthesia.

##### Crude estimates of relevant anaesthetic drug concentrations based on simplifying assumptions

Although it is difficult to compare the relevant drug concentrations *in vitro* vs *in vivo*, here we make crude estimates based on the following simplifying assumptions. We assume, for both ketamine and propofol: (A) that essentially all the drug added *in vitro* is free (i.e. in aqueous solution) in the aCSF, i.e. not lost to degradation or binding to vials, tubing etc.; (B) that there is full equilibration *in vitro*, between the free (in aqueous solution) drug concentrations in the aCSF that we used to perfuse our slice bath and the free drug concentrations at the site of action in neurons within the slices after 2 h of exposure;; and also (C) full equilibration *in vivo*, between free drug concentrations in the blood and at the site of action in neurons within the brain. Although we use these simple assumptions for making initial, crude estimates, they are oversimplifications that can lead to substantial errors, in particular for *in vitro* experiments (see below).

#### Ketamine

##### Crude estimates of ketamine concentrations based on human data

For ketamine, we estimate, based on the above assumptions and data from humans: (1) The 20 µM ketamine used in our aCSF for slice experiments (**f**) corresponds roughly to (i.e. should give ~the same local concentration at the site of action in neurons as) a total blood concentration of 40 µM ketamine in humans, because about half of the total ketamine in blood is bound to protein (~47%; (Dayton et al. 1983); giving ~20 µM free ketamine; 20 µM = 40µM/2). (2) Studies in humans have estimated that full, general anaesthesia (unresponsiveness to both verbal command and pain) caused by ketamine alone requires at least a total ketamine concentration in the blood (50% free plus 50% protein bound) of about 3-4 µM (Schüttler et al. 1987), yielding ~2µM (~4µM/2) free ketamine in blood. From (1) and (2) we conclude that our in vitro concentration of 20 µM free ketamine is about 10 times higher (20 µM vs. 2 µM) than the *in vivo* concentration of free ketamine in the human brain required for full surgical anaesthesia with ketamine alone. This estimate should hold if our simplifying assumptions hold; but as discussed below, there are complicating factors.

##### Ketamine concentrations *in vitro* vs. *in vivo*, in rodents

In previous *in vitro* experiments with ketamine in rodent CNS slices, a wide range of concentrations have been used, from 30 nM to 10 mM free ketamine (Schnoebel et al. 2005; Yin et al. 2019). *In vivo*, in rodents, intraperitoneal (IP) injections of sub-anaesthetic ketamine doses (7.5-25 mg/kg) yielded total ketamine concentrations in brain tissue ranging from 4 µM to 11 µM (Zanos et al. 2016; Yang et al. 2018). This should correspond to free ketamine concentrations in brain tissue that are higher than 1.3-3.6 µM, since the total ketamine concentration in brain slices is about 3-fold higher than the free ketamine in the surrounding aCSF after prolonged equilibration e.g. 50 µM total vs. 17 µM free (Geiger et al. 2021). This reflects that most of the total ketamine in brain tissue is bound after long equilibration, but the ratio between total and free ketamine must be lower after briefer exposures that do not allow ketamine to fully bind in the tissue.

##### Effects of ketamine *in vitro* vs. *in vivo*, in rodents

We saw no significant effect on PPs of relatively low ketamine (20 µM), neither with wash-in nor pre-incubation (**Figures 2** and **3**). However, if the reported sub-anaesthetic brain concentrations in rodents are accurate (4-11 µM total ketamine in brain after IP injections of sub-anaesthetic doses: 7.5-25 mg/kg (Zanos et al. 2016; Yang et al. 2018)), it seems likely that supra-anaesthetic doses give ~10 fold higher brain concentrations, since reported anaesthetic IP doses in rodents are 100-190 mg/kg (~10-fold higher than sub-anaesthetic doses 7.5-25 mg/kg); this yields full anaesthesia in 100% of animals (Levin-Arama et al. 2016; Cichon et al. 2022). This suggests a free brain concentration of ketamine during full anaesthesia of (4-11 µM x 10 =) 40-110 µM, resembling the 20-100 µM concentrations that we used in the aCSF for our wash-in and pre-incubation experiments (**Figures 2** and **3**).
It may, however, seem unclear why this estimate (20-100 µM) is ~5-25 times higher than our above crude estimate (based on simplifying assumptions), which was 4 µM free ketamine in the human brain during full ketamine anaesthesia. One possibility is that rodents may require higher free brain concentration of ketamine than humans for full anaesthesia (Ribeiro et al. 2014) or the estimates may be inaccurate. It should be noted that the IV starting dose to induce general anaesthesia in humans is typically 3-4 mg/kg (Idvall et al. 1979) whereas the above-mentioned IP doses in rodents are in the 100-190 mg/kg range. Furthermore, the above wide ranges (20-100 µM; ~5-25 times higher than our initial estimate) may reflect that it is very difficult to determine the exact minimal dose of ketamine for full anaesthesia caused by ketamine alone in rodents; hence some reported values may actually be well above that minimum. Alternatively, the apparent difference in anaesthetic dose between rodents and humans may be mainly caused by a difference between intravenous (IV) vs. intraperitoneal (IP) injections. Thus, IV injections seem to be much more effective than IP injections of similar ketamine doses in rodents: IV bolus injections of 20-30 mg/kg yielded measured total brain concentrations of up to 300-360 µM (Cohen et al. 1973; Cohen and Trevor 1974;

Marietta et al. 1976), whereas IP injections of comparable ketamine doses (7.5-25 mg/kg) yielded far lower total ketamine concentrations in the rodent brains: 4-11 µM (Zanos et al. 2016; Yang et al. 2018). Also, in paediatric anaesthesia, IP doses are 4-5 times higher than IV doses (Grant et al. 1983). This IV vs. IP difference may be caused by slow and incomplete absorption into the blood when IP injections are used.

##### Possible relevance of our *in vitro* results with ketamine to anaesthetic and other effects *in vivo*

These estimates, based on *in vivo* ketamine anaesthesia in rodents (20-100 µM free ketamine in the brain) suggest that the different effects on plateau potentials (PPs) that we found between these concentrations (100 µM ketamine reduced the PPs, whereas 20 µM had little or no effects on the PPs: **Figures 2** and **3**), may be relevant for the *in vivo* anaesthetic effects of ketamine, at least in rodents. Furthermore, this reasoning, based on rodent data, may suggest that the plateau suppression that we saw at higher but not lower concentrations of ketamine (**Figures 2** and **3**) may contribute to the suppression of responsiveness combined with altered “inner” consciousness seen at anaesthetic doses in humans (Sarasso et al. 2015).

##### Apparent bimodal concentration-dependence of ketamine effects on PPs: possible relevance for consciousness?

We found that the mean values of the PP area and spike frequency were somewhat larger after preincubation with 20 µM ketamine than without ketamine (**Figure 3 Ci-ii**). This might suggest that relatively low ketamine concentrations increased the plateaus whereas higher concentrations reduced the plateaus, thus suggesting a bimodal, concentration-dependent ketamine effect. We do not know whether this surprising pattern is real or due to chance, or might reflect different subpopulations of L2/3PCs that we could not distinguish due to small sample sizes. However, if there is really such a bimodal concentration-dependence of the ketamine effect on the PPs, it seems to fit well with, and may help explain, certain observations of seemingly bimodal, concentration-dependent effects of ketamine *in vivo* both in humans (Farnes et al. 2020) and in rodents (Arena et al. 2022). It may correspond to the fact that the human EEG shows a pattern of arousal and high activity during clinical ketamine anaesthesia, very different from the general EEG suppression seen with propofol and most other anaesthetic agents (Akeju et al. 2016, 2014).

The apparent enhancing effect on PPs of relatively low ketamine (20 µM) in both wash-in and pre-incubation experiments (**Figures 2** and **3**), may suggest that low ketamine doses may exert their potential antidepressant effects (Mion and Villevieille 2013) by enhancing PPs and persistent firing. This possibility may be supported by evidence suggesting that sub-anaesthetic, antidepressant doses of ketamine cause a hyperglutamatergic state in the PFC, in contrast to anaesthetic doses which decrease PFC glutamate levels (Moghaddam et al. 1997). This again suggests a bimodal concentration-dependence, which might be related to an enhanced PP-driven firing and glutamate release induced by low ketamine (**Figure 3 Ci-ii**).

#### Propofol

##### Crude estimates of propofol concentrations based on human data

For propofol, we can make at last two estimates, based on the above assumptions and data from humans:

Estimate 1: (1) Measured total blood plasma concentrations of propofol (including protein-bound) during surgical general anaesthesia range from ~3 to 27 mg/ml (Braathen et al. 2022; Smith et al. 1994), which converts to ~16-153 µM. However, it is important to note that the large range reported by Smith et al. (1994) reflects very different forms of "unresponsiveness": suppression of responses to speech in 95% of subjects require a blood concentration of only 5.4 µg/ml propofol , whereas suppression of responses to painful surgery (incisions) in 95% of the same subjects, 27.4 propofol µg/ml was required. Since propofol has been reported to be ~97% bound to plasma proteins (Cockshott et al. 1992), the above total plasma concentrations (~16-153 µM), should yield ~0.5-4.6 µM free, unbound propofol. This suggests that the 3 µM propofol used in our aCSF for slice experiments (**Figures 4** and **5**) is close to the concentration range relevant for deep general anaesthesia and analgesia with propofol only in humans. However, this dosing schedule which is almost never used for surgery, as propofol is used in the 3-4 µg/ml total plasma concentration range (corresponding to ~0.5 µM free, unbound propofol) for verbal unconsciousness, supplemented with an analgesic drug (i.e. opioids or other) for analgesic unresponsiveness,but other estimates support different conclusions.

Estimate 2: (1) Other studies in humans have estimated that full, general anaesthesia (unresponsiveness) caused by propofol together with adjuvants requires a total propofol concentration in the blood of about 3-5 µM (Braathen et al. 2022) or 6 µM (35 µg/ml blood; Struys et al. 2000). Assuming ~95% protein binding of propofol in blood as described by Ishii-Maruhama et al. (2018), 5 µM total concentration yields ~0.25 µM (5% of 5 µM) free propofol in blood during anaesthesia. (2) However, the 3 µM propofol that we used in our aCSF for slice experiments (**Figures 4** and **5**) corresponds roughly to (i.e. should give ~the same local concentration at the site of action in neurons as) a total blood concentration of 60 µM propofol, assuming ~95% protein binding in blood (giving ~3 µM free propofol, i.e. 5% of 60 µM). From (1) and (2) we conclude that our *in vitro* concentration of 3 µM free propofol is about 10 times higher than the *in vivo* concentration (~0.25 µM) of free propofol in the human brain required for adequate clinical propofol anaesthesia when combined with adjuvants. Again, this estimate holds only if both the reported data and our simplifying assumptions hold, but this is unlikely, as there are again several complicating factors, as discussed below.

The fact that our two estimates (1 and 2) differ substantially, may support the conclusion that such simplified estimates are not accurate.Thus, for propofol the estimates will vary 4-5-fold depending on whether propofol is used for painful surgery as mono-anaesthetic (Estimate 1) or (as in clinical anaesthesia) together with analgesic adjuvants. Also contributing to the discrepancy may be different criteria being used for defining "unresponsiveness" in humans and animals. In general, evidence suggests that *in vitro* experiments often require higher drug concentrations than *in vivo*, but the mechanisms are complex and often poorly understood (Leist et al. 2017; Hengstler et al. 2020).

#### Long pre-incubation of propofol was needed to reveal effects of lower doses

A major difference between intravenous administration of ketamine or propofol *in vivo* and our bath-application of these drugs *in vitro*, is that brain slices of course lack the *in vivo* blood supply through a dense capillary net, which brings drugs rapidly to within a few micrometres from each brain cell, requiring minimal diffusion time. This difference proved to be particularly important for propofol and probably explains its far slower equilibration in slices.

In line with this reasoning, our experiments using 2 hours of pre-incubation with relatively low concentrations of propofol (3 µM) or ketamine (20 µM) showed considerably stronger drug effects than our wash-in experiments applying the same drug concentrations for 5-25 minutes. This difference was particularly striking for propofol (compare **Figures 2** and **3** for ketamine, and **Figures 4** and **5** for propofol). This strongly suggests that our wash-in applications did not reveal the full effect of the applied drug concentrations, presumably because of delayed equilibration between the applied medium and the interior of the slice. Although this interpretation may seem surprising in view of the fast anaesthetic effects of these drugs when given IV *in vivo* (Scheepstra et al. 1989; Baekgaard et al. 2019), it is strongly supported by recent studies of *in vitro* diffusion of ketamine and propofol (Gredell et al. 2004; Geiger et al. 2021) and thus probably correct. Therefore, we base our discussion and conclusions primarily on our 2 hour pre-incubation results.

#### Effects of ketamine and propofol on PPs vs. states of consciousness

How may the similarity in results between high concentrations of ketamine and propofol at higher concentrations (100 µM and 10 µM respectively, **Figures 2** and **4**) be related to their dramatically different states of unconsciousness? The similarity between the effects of ketamine and propofol that we observed suggests that PPs may have a greater role in supporting external awareness and responsiveness than on conscious experience itself, as reportable “inner” consciousness in the form of vivid dreaming is preserved in ketamine but less so in propofol anaesthesia whilst responsiveness are abolished in both (Sarasso et al. 2015; Nilsen et al. 2022).

#### Statistics and cell types

We also want to address the fact that several cells out of our sample lie outside the 95% confidence interval (CI; see **Figures 2C** and **4C**). As shown in several studies, there are subtypes of L2/3PCs and other cortical pyramidal cell types in rodents (van Aerde and Feldmeyer 2015; Yao et al. 2023). Perhaps the great spread in response reflects the diversity among subtypes of cells, as this pattern was seen in our study across three different patch clamp rigs and experimentalists.

#### Other sources of error for *in vitro* propofol experiments

In our early, preliminary experiments with propofol, we were initially unable to replicate effects on electrophysiological properties reported in the literature such as inhibition of the $I_h$ (HCN) current (Ying et al. 2006). Previous studies have indicated absorption of propofol by silicon and Tygon tubing (Maurer et al. 2017), which we initially used to circulate aCSF to our recording chamber, and also absorption of propofol to certain types of plastics during storage (Sautou-Miranda et al. 1996). After we changed the tubing, vials, and drug preparation, we found reliable effects of propofol (**Figures 2** and **3**). In addition, bacterial degradation of the drugs (in standard, non-sterile *in vitro* conditions) may also reduce their effective concentrations over time. These findings suggest that also the first (A) of our "simplifying assumptions", used for our initial, crude estimates (see above), do not always hold.

The interactions between propofol and materials commonly used in *in vitro* experiments (Maurer et al. 2017; Sautou-Miranda et al. 1996), may call into question the qualitative and quantitative effects of propofol previously reported in some slice studies, as absorption of propofol may lead to overestimation of the concentration required to achieve particular effects.

#### References

Aerde, Karlijn I. van, and Dirk Feldmeyer. 2015. "Morphological and Physiological Characterization of Pyramidal Neuron Subtypes in Rat Medial Prefrontal Cortex." Cerebral Cortex 25 (3): 788–805.

Akeju, Oluwaseun, Andrew H. Song, Allison E. Hamilos, Kara J. Pavone, Francisco J. Flores, Emery N. Brown, and Patrick L. Purdon. 2016. "Electroencephalogram Signatures of Ketamine Anesthesia-Induced Unconsciousness." Clinical Neurophysiology: Official Journal of the International Federation of Clinical Neurophysiology 127 (6): 2414–22.

Akeju, Oluwaseun, M. Brandon Westover, Kara J. Pavone, Aaron L. Sampson, Katharine E. Hartnack, Emery N. Brown, and Patrick L. Purdon. 2014. "Effects of Sevoflurane and Propofol on Frontal Electroencephalogram Power and Coherence." Anesthesiology 121 (5): 990–98.

Arena, A., B. E. Juel, R. Comolatti, S. Thon, and J. F. Storm. 2022. "Capacity for Consciousness under Ketamine Anaesthesia Is Selectively Associated with Activity in Posteromedial Cortex in Rats." Neuroscience of Consciousness 2022 (1): niac004.

Baekgaard, Josefine S., Trine G. Eskesen, Martin Sillesen, Lars S. Rasmussen, and Jacob Steinmetz. 2019. "Ketamine as a Rapid Sequence Induction Agent in the Trauma Population: A Systematic Review." Anesthesia and Analgesia 128 (3): 504–10.

Braathen, Martin R., Ivan Rimstad, Terje Dybvik, Ståle Nygård, and Johan Raeder. 2022. "Online Exhaled Propofol Monitoring in Normal-Weight and Obese Surgical Patients." Acta Anaesthesiologica Scandinavica 66 (5): 598–605.

Braithwaite, Hannah E., Thomas Payne, Nicholas Duce, Jessica Lim, Tim McCulloch, John Loadsman, Kate Leslie, Angela C. Webster, Amy Gaskell, and Robert D. Sanders. 2023. "Impact of Female Sex on Anaesthetic Awareness, Depth, and Emergence: A Systematic Review and Meta-Analysis." British Journal of Anaesthesia 131 (3): 510–22.

Cichon, Joseph, Andrzej Z. Wasilczuk, Loren L. Looger, Diego Contreras, Max B. Kelz, and Alex Proekt. 2022. "Ketamine Triggers a Switch in Excitatory Neuronal Activity across Neocortex." Nature Neuroscience 26 (1): 39–52.

Cockshott, I. D., E. J. Douglas, G. F. Plummer, and P. J. Simons. 1992. "The Pharmacokinetics of Propofol in Laboratory Animals." Xenobiotica; the Fate of Foreign Compounds in Biological Systems 22 (3): 369–75.

Cohen, M. L., S. L. Chan, W. L. Way, and A. J. Trevor. 1973. "Distribution in the Brain and Metabolism of Ketamine in the Rat after Intravenous Administration." Anesthesiology 39 (4): 370–75.

Cohen, M. L., and A. J. Trevor. 1974. "On the Cerebral Accumulation of Ketamine and the Relationship between Metabolism of the Drug and Its Pharmacological Effects." The Journal of Pharmacology and Experimental Therapeutics 189 (2): 351–58.

Dayton, P. G., R. L. Stiller, D. R. Cook, and J. M. Perel. 1983. "The Binding of Ketamine to

Plasma Proteins: Emphasis on Human Plasma." European Journal of Clinical Pharmacology 24 (6): 825–31.
Farnes, Nadine, Bjørn E. Juel, André S. Nilsen, Luis G. Romundstad, and Johan F. Storm. 2020. "Increased Signal Diversity/complexity of Spontaneous EEG, but Not Evoked EEG Responses, in Ketamine-Induced Psychedelic State in Humans." PloS One 15 (11): e0242056.
Franks, N. P., and W. R. Lieb. 1994. "Molecular and Cellular Mechanisms of General Anaesthesia." Nature 367 (6464): 607–14.
Geiger, Zachary, Brett VanVeller, Zarin Lopez, Abdel K. Harrata, Kathryn Battani, Lauren Wegman-Points, and Li-Lian Yuan. 2021. "Determination of Diffusion Kinetics of Ketamine in Brain Tissue: Implications for in Vitro Mechanistic Studies of Drug Actions." Frontiers in Neuroscience. https://doi.org/10.3389/fnins.2021.678978.
Grant, I. S., W. S. Nimmo, L. R. McNicol, and J. A. Clements. 1983. "Ketamine Disposition in Children and Adults." British Journal of Anaesthesia 55 (11): 1107–11.
Gredell, J. A., P. A. Turnquist, M. B. Maciver, and R. A. Pearce. 2004. "Determination of Diffusion and Partition Coefficients of Propofol in Rat Brain Tissue: Implications for Studies of Drug Action in Vitro." British Journal of Anaesthesia 93 (6): 810–17.
Hengstler, Jan G., Anna-Karin Sjögren, Daniele Zink, and Jorrit J. Hornberg. 2020. "In Vitro Prediction of Organ Toxicity: The Challenges of Scaling and Secondary Mechanisms of Toxicity." Archives of Toxicology 94 (2): 353–56.
Hijazi, Youssef, and Roselyne Boulieu. 2002. "Protein Binding of Ketamine and Its Active Metabolites to Human Serum." European Journal of Clinical Pharmacology 58 (1): 37–40.
Idvall, J., I. Ahlgren, K. R. Aronsen, and P. Stenberg. 1979. "Ketamine Infusions: Pharmacokinetics and Clinical Effects." British Journal of Anaesthesia 51 (12): 1167–73.
Ishii-Maruhama, Minako, Hitoshi Higuchi, Mai Nakanou, Yuka Honda-Wakasugi, Akiko Yabuki-Kawase, Shigeru Maeda, and Takuya Miyawaki. 2018. "In Vitro Changes in the Proportion of Protein-Unbound-Free Propofol Induced by Valproate." Journal of Anesthesia 32 (5): 688–93.
Leist, Marcel, Ahmed Ghallab, Rabea Graepel, Rosemarie Marchan, Reham Hassan, Susanne Hougaard Bennekou, Alice Limonciel, et al. 2017. "Adverse Outcome Pathways: Opportunities, Limitations and Open Questions." Archives of Toxicology 91 (11): 3477–3505.
Levin-Arama, Maya, Lital Abraham, Trevor Waner, Alon Harmelin, David M. Steinberg, Tal Lahav, and Mickey Harlev. 2016. "Subcutaneous Compared with Intraperitoneal KetamineXylazine for Anesthesia of Mice." Journal of the American Association for Laboratory Animal Science: JAALAS 55 (6): 794–800.
Marietta, M. P., P. F. White, C. R. Pudwill, W. L. Way, and A. J. Trevor. 1976. "Biodisposition of Ketamine in the Rat: Self-Induction of Metabolism." The Journal of Pharmacology and Experimental Therapeutics 196 (3): 536–44.
Maurer, F., D. J. Lorenz, G. Pielsticker, T. Volk, D. I. Sessler, J. I. Baumbach, and S. Kreuer. 2017. "Adherence of Volatile Propofol to Various Types of Plastic Tubing." Journal of Breath Research 11 (1): 016009.
Mion, Georges, and Thierry Villevieille. 2013. "Ketamine Pharmacology: An Update (Pharmacodynamics and Molecular Aspects, Recent Findings)." CNS Neuroscience & Therapeutics 19 (6): 370–80.
Moghaddam, B., B. Adams, A. Verma, and D. Daly. 1997. "Activation of Glutamatergic Neurotransmission by Ketamine: A Novel Step in the Pathway from NMDA Receptor Blockade to Dopaminergic and Cognitive Disruptions Associated with the Prefrontal Cortex." The Journal of Neuroscience: The Official Journal of the Society for Neuroscience 17 (8): 2921–27.
Nilsen, André S., Bjørn E. Juel, Benjamin Thürer, Arnfinn Aamodt, and Johan F. Storm. 2022. "Are We Really Unconscious in 'unconscious' States? Common Assumptions

Revisited.” Frontiers in Human Neuroscience 16 (October): 987051.
Pai, A., and M. Heining. 2007. “Ketamine.” Continuing Education in Anaesthesia Critical Care & Pain 7 (2): 59–63.
Ribeiro, Patrícia O., Angelo R. Tomé, Henrique B. Silva, Rodrigo A. Cunha, and Luís M. Antunes. 2014. “Clinically Relevant Concentrations of Ketamine Mainly Affect Long-Term Potentiation rather than Basal Excitatory Synaptic Transmission and Do Not Change Paired-Pulse Facilitation in Mouse Hippocampal Slices.” Brain Research 1560 (April): 10–17.
Sarasso, Simone, Melanie Boly, Martino Napolitani, Olivia Gosseries, Vanessa Charland-Verville, Silvia Casarotto, Mario Rosanova, et al. 2015. “Consciousness and Complexity during Unresponsiveness Induced by Propofol, Xenon, and Ketamine.” Current Biology: CB 25 (23): 3099–3105.
Sautou-Miranda, V., E. Levadoux, M. T. Groueix, and J. Chopineau. 1996. “Compatibility of Propofol Diluted in 5% Glucose with Glass and Plastics (polypropylene, Polyvinylchloride) Containers.” International Journal of Pharmaceutics 130 (2): 251–55.
Scheepstra, G. L., L. H. Booij, C. L. Rutten, and L. G. Coenen. 1989. “Propofol for Induction and Maintenance of Anaesthesia: Comparison between Younger and Older Patients.” British Journal of Anaesthesia 62 (1): 54–60.
Schnoebel, Rose, Matthias Wolff, Saskia C. Peters, Michael E. Bräu, Andreas Scholz, Gunter Hempelmann, Horst Olschewski, and Andrea Olschewski. 2005. “Ketamine Impairs Excitability in Superficial Dorsal Horn Neurones by Blocking Sodium and Voltage-Gated Potassium Currents.” British Journal of Pharmacology 146 (6): 826–33.
Schüttler, J., D. R. Stanski, P. F. White, A. J. Trevor, Y. Horai, D. Verotta, and L. B. Sheiner. 1987. “Pharmacodynamic Modeling of the EEG Effects of Ketamine and Its Enantiomers in Man.” Journal of Pharmacokinetics and Biopharmaceutics 15 (3): 241–53.
Searle, R., and P. M. Hopkins. 2009. “Pharmacogenomic Variability and Anaesthesia.” British Journal of Anaesthesia. https://doi.org/10.1093/bja/aep130.
Smith, C., A. I. McEwan, R. Jhaveri, M. Wilkinson, D. Goodman, L. R. Smith, A. T. Canada, and P. S. Glass. 1994. “The Interaction of Fentanyl on the Cp50 of Propofol for Loss of Consciousness and Skin Incision.” Anesthesiology 81 (4): 820–28; discussion 26A.
Struys, M. M., T. De Smet, B. Depoorter, L. F. Versichelen, E. P. Mortier, F. J. Dumortier, S. L. Shafer, and G. Rolly. 2000. “Comparison of Plasma Compartment versus Two Methods for Effect Compartment--Controlled Target-Controlled Infusion for Propofol.” Anesthesiology 92 (2): 399–406.
Yang, Yan, Yihui Cui, Kangning Sang, Yiyan Dong, Zheyi Ni, Shuangshuang Ma, and Hailan Hu. 2018. “Ketamine Blocks Bursting in the Lateral Habenula to Rapidly Relieve Depression.” Nature 554 (7692): 317–22.
Yao, Zizhen, Cindy T. J. van Velthoven, Michael Kunst, Meng Zhang, Delissa McMillen, Changkyu Lee, Won Jung, et al. 2023. “A High-Resolution Transcriptomic and Spatial Atlas of Cell Types in the Whole Mouse Brain.” Nature 624 (7991): 317–32.
Ying, Shui-Wang, Syed Y. Abbas, Neil L. Harrison, and Peter A. Goldstein. 2006. “Propofol Block of I(h) Contributes to the Suppression of Neuronal Excitability and Rhythmic Burst Firing in Thalamocortical Neurons.” The European Journal of Neuroscience 23 (2): 465–80.
Yin, Jianyin, Bao Fu, Yuan Wang, and Tian Yu. 2019. “Effects of Ketamine on Voltage-Gated Sodium Channels in the Barrel Cortex and the Ventral Posteromedial Nucleus Slices of Rats.” Neuroreport 30 (17): 1197–1204.
Zanos, Panos, Ruin Moaddel, Patrick J. Morris, Polymnia Georgiou, Jonathan Fischell, Greg I. Elmer, Manickavasagom Alkondon, et al. 2016. “NMDAR Inhibition-Independent Antidepressant Actions of Ketamine Metabolites.” Nature 533 (7604): 481–86.
